# Loss of Lamp2a-dependent chaperone-mediated autophagy drives dry AMD-like retinal pathology in mice and is rescued by BK channel activation

**DOI:** 10.64898/2026.03.19.712761

**Authors:** Hilal Ahmad Mir, Gaddam Mahesh, Arunan Palanimuthu, Christopher L. Cioffi, Konstantin Petrukhin

**Affiliations:** Department of Ophthalmology, Columbia University Irving Medical Center, Columbia University, New York, NY 10032, USA; Rensselaer Polytechnic Institute, 110 8th Street, Troy, New York 12180, USA

**Keywords:** Dry age-related macular degeneration, Chaperone-mediated autophagy, Lamp2a, Retinal pigment epithelium, BK channel activation, Autophagic flux

## Abstract

Age-related macular degeneration (AMD) is the leading cause of irreversible visual loss in elderly individuals for which no effective treatments are currently available. The photoreceptor loss in dry AMD is secondary to the demise of the retinal pigment epithelium (RPE) cells. The accumulation of extracellular deposits, known as drusen, resulting in part from deficient lysosomal and autophagosomal degradation, is a key feature of dry AMD pathogenesis. Chaperone-mediated autophagy (CMA) is a selective lysosomal degradation pathway that maintains proteostasis by targeting specific cytosolic proteins for lysosomal translocation and degradation. LAMP2A (lysosome-associated membrane protein 2A) functions as the key lysosomal receptor required for CMA. Using *Lamp2a* knockout mouse, we show that selective CMA dysfunction recapitulates AMD-like pathologies, including sub-RPE lipid and protein deposits, RPE atrophy, Bruch’s membrane thickening, and impaired autophagic activity. Furthermore, we identify large-conductance Ca²⁺-activated K⁺ (BK) channels as a therapeutic target for restoring autophagic activity. Mechanistically, pharmacological activation of BK channels with the small-molecule agonist GLA-1-1 enhances macroautophagy and stimulates autophagic flux by promoting autophagosome-lysosome fusion. Importantly, oral administration of GLA-1-1 in markedly attenuates structural, functional, and molecular retinal abnormalities in *Lamp2a*-deficient mice, suggesting that pharmacological activation of macroautophagy through facilitating autophagosome-lysosome fusion can partially compensate for CMA deficiency. Together, these findings demonstrate that pharmacological activation of macroautophagy can ameliorate the retinal phenotype resulting from CMA dysfunction and support BK channel activation by GLA-1-1 as a promising therapeutic strategy for dry AMD.

## Introduction

Age related macular degeneration (AMD) is the most common cause of central vison loss in adults over 60 globally. AMD exists in two forms wet or neovascular form (15% of cases) and dry or atrophic form (85% of cases) [1]. Approved anti-complement therapies for dry AMD (Syfovre and Izervay) require frequent intraocular injections, offer limited efficacy, and have been reported in some studies to potentially increase the risk of developing the more severe wet form of AMD [2–5]. Dry AMD is a progressive disease that begins with the accumulation of lipoprotein-rich extracellular deposits known as drusen during the early and intermediate stages [6]. Soft sub-RPE drusen are an established risk factor for progression to advanced disease. At the late stage, degeneration of the RPE and loss of photoreceptors in the retina manifest as areas of geographic atrophy (GA) [7]. Besides, sub-RPE drusen, formation of another type of extracellular deposits, reticular pseudodrusen (RPD) located over the apical side of the RPE [8], is strongly associated with GA growth [9]. Although the mechanisms of drusen biogenesis are not completely known, several studies have linked impaired clearance of aggregated proteins by the lysosome-autophagosome and lysosome-phagosome networks in the RPE as a primary factor contributing to drusen accumulation [10]. RPE cells play an important role in maintaining retinal health by digesting photoreceptor outer segments (POS) via phagocytosis. Incomplete POS digestion generates unwanted material that contributes to drusen formation [11]. Autophagy and phagocytosis are interdependent and share common maturation steps; both rely on efficient fusion of autophagosomes and phagosomes with lysosomes for proper cargo degradation [12]. Lysosomes are at the crossroads of the two degradation pathways that are critical in biogenesis of drusen: phagocytosis and autophagy [13]. Defective fusion of lysosomes with phagosomes and autophagosomes relate to a variety of neurodegenerative diseases characterized by misfolding, aggregation and accumulation of protein deposits [14].

Autophagy comprises three lysosome-dependent catabolic pathways that mediate intracellular degradation: macroautophagy, microautophagy, and chaperone-mediated autophagy (CMA). Recently, deficient autophagy has been shown to produce an AMD-like phenotype in a genetic mouse model: mice lacking lysosome-associated membrane protein type 2 gene, *Lamp2*, showed accumulation of basal laminar deposits in the retina and accumulation of drusen associated proteins [15]. Patients with Danon disease, caused by loss-of-function mutations in *LAMP2*, exhibit retinal degeneration and maculopathy as part of the disease phenotype.[16–19] Both Danon disease and *Lamp2* knockout mice are associated with high mortality. LAMP2 is lysosomal glycoprotein that plays a critical role in lysosomal biogenesis and function. There are three LAMP2 isoforms generated by alternative splicing of exon 9 of the *LAMP2* gene: LAMP2A, which functions as the receptor and channel for chaperone-mediated autophagy (CMA), LAMP2B, which is required for autophagosome-lysosome fusion, and LAMP2C, which mediates lysosomal uptake and degradation of nucleic acids [20]. CMA is a selective degradative pathway in which cytosolic proteins containing a KFERQ-like motif are recognized by molecular chaperones and translocated across the lysosomal membrane through interaction with the LAMP2A receptor [21]. CMA mediated autophagy declines with age and has been linked to several neurodegenerative diseases including progressive retinal degeneration, suggesting the potential role of LAMP2A in retinal homeostasis [22–24]. It has been shown that CMA activity is markedly impaired in AMD patients, and that iPSC-derived RPE from these patients exhibit significantly reduced LAMP2A expression that disrupts CMA-mediated proteostasis [25]. Considering that genetic ablation of all Lamp2 isoforms produces an AMD-like phenotype but is associated with high mortality [15], that CMA activity is markedly impaired in AMD patients [25], and that Lamp2a may be the predominant LAMP2 isoform expressed in the RPE, we studied *Lamp2a*^-/-^ mice to additionally investigate the role of CMA in processes relevant to dry AMD pathogenesis, with the expectation that selective deletion of *Lamp2a* might be associated with reduced mortality.

In parallel, we sought to explore potential therapeutic interventions aimed at restoring lysosomal and autophagic function in dry AMD. BK (big potassium) channels, also known as maxi-K^+^, Slo1, and KCa1.1 channels, are characterized by large K^+^ conductance, sensitivity to voltage and Ca^2+^, and ubiquitous expression.[26] Plasma membrane BK channels play critical roles in regulating neuronal excitability, neurotransmitter release, endocrine secretion, muscle contractility, epithelial function, and other functions.[27] BK channels were shown to be present in lysosomes of most cell types. [28, 29] The role of BK channels in lysosomal function relates to regulation of the Ca^2+^ release from lysosomes[28, 30] and to Ca^2+^ refilling into lysosomes following its depletion.[29] A functional coupling between BK and Ca^2+^ channels exists in lysosomes.[28, 30] Activation of the lysosomal BK channels provides a positive feedback regulation in which K^+^ entry into the lysosomal lumen provides a counter-ion and a driving force for the Ca^2+^ efflux to the cytoplasm.[28, 30] The net result is the increase in Ca^2+^ release through lysosomal TRML1[28, 30], CACNA1A[31], and other types of Ca channels.[32] Ca^2+^ release from the fusing organelles (such as autophagosomes and lysosomes or phagosomes and lysosomes) is required for the fusing event and effective lysosomal degradation.[33] Induction of the Ca^2+^ release from lysosomes is shown to facilitate the SNARE-mediated fusion of lysosomes with autophagosomes and improves degradation of the autophagosomal cargo.[31] BK channel overexpression[28] or its pharmacological activation with agonists [30] stimulates the lysosomal Ca^2+^ release and rescues the disease phenotypes associated with the deficiency in lysosomal cargo degradation and trafficking [28, 30]. Boosting the activity of lysosomal TRPML1, a functional partner of BK, with an agonist increases the Ca^2+^ release and restores the autophagic flux in cells from Lowe syndrome patients.[34] Loss of BK channel activity causes dysfunction in lysosomal trafficking and cargo degradation.[28, 29] Pharmacological inhibition of the BK channel suppresses phagocytosis.[35] Based on available evidence, we hypothesize that pharmacological activation of lysosomal BK channels may optimize autophagosome-lysosome (and phagosome-lysosome) fusion, thereby normalizing lysosomal degradation of autophagic and phagocytic cargo, which may be relevant for inhibiting drusen formation in dry AMD. While CMA, which appears to play an important role in mediating pathogenic changes in the RPE, is a receptor-mediated pathway that does not require autophagosome-lysosome fusion, CMA deficiency resulting from LAMP2A abnormalities may be partially compensated by increased macroautophagy, which involves fusion of autophagosomes with lysosomes, as previously suggested. [36, 37]. In the current study, we show that autophagic flux and autophagosome-lysosome fusion in cultured cells are inhibited by the selective BK channel inhibitor penitrem A, indicating BK channel dependence of this process. Conversely, the small-molecule BK channel opener GLA-1-1 induces autophagic flux and promotes autophagosome-lysosome fusion. We further demonstrate that *Lamp2a⁻^/^⁻*mice recapitulate key features of dry AMD, including accumulation of autofluorescent granules, Bruch’s membrane thickening, accumulation of drusen-associated proteins and lipid droplets, and increased autophagosome abundance. Remarkably, these pathological changes were significantly attenuated in *Lamp2a* knockout mice fed GLA-1-1-formulated chow. Collectively, these findings highlight a pivotal contribution of CMA dysfunction to dry AMD pathogenesis and identify BK channel activation by GLA-1-1 as a potential therapeutic strategy to restore autophagic flux and mitigate pathogenic processes involved in dry AMD

## Results

### 1. GLA-1-1 induces potassium channel dependent autophagy flux by promoting autophagosome-lysosome fusion

Over the years, activation of plasma membrane BK channels has been proposed as a therapeutic strategy for smooth muscle disorders such as hypertension, ischemic heart disease, airway hyperexcitability, urinary incontinence, stroke, and erectile dysfunction. [38, 39] A variety of small-molecule BK channel activators from different structural classes have been reported. [38, 40, 41] However, none have been approved by regulatory agencies, and clinical evaluation of BK agonists has largely been discontinued. [38] One of the most active and best characterized BK agonists is GLA-1-1 (Fig. 1B, C; Fig. S1A, B), originally described in efforts to develop therapies for smooth muscle disorders.[42] In our hands, GLA-1-1 was more potent than other known BK agonists (NS1619, NS11021, and MaxiPost) in a previously described FLIPR assay [42], with an EC₅₀ of 3.4 ± 2.1 µM. The agonistic activity observed in the FLIPR assay was abolished by the selective BK inhibitor penitrem A. Patch-clamp electrophysiology confirmed activation of BK channels by GLA-1-1 (Fig. 1D; Fig. S1C). Based on its robust BK channel agonistic activity, GLA-1-1 was used to probe the role of BK channel activation in autophagy. Fusion of autophagosomes with lysosomes is a key step in autophagic flux. Activation of lysosomal BK channels promotes potassium influx that supports lysosomal calcium release, thereby facilitating autophagosome-lysosome fusion. We investigated whether BK channel activation by GLA-1-1 could facilitate autophagosome-lysosome fusion and modulate autophagic flux. To facilitate quantitative assessment of autophagic flux, stable HeLa reporter clones expressing tandem mCherry-EGFP-LC3B were generated by retroviral transduction followed by selection and fluorescence-activated cell sorting (FACS) of mCherry-GFP double-positive cells (Fig. S1D). This reporter cell line enables quantitative assessment of autophagic flux based on the differential pH sensitivity of GFP and mCherry, whereby acidification of LC3-positive vesicles following autophagosome-lysosome fusion quenches GFP fluorescence while mCherry signal is retained. Consequently, increased autophagosome-lysosome fusion and autophagic flux are reflected by an increased cellular mCherry/GFP fluorescence ratio. FACS analysis showed that serum starvation markedly increased the mCherry/GFP ratio, an effect abolished by the BK channel inhibitor penitrem A, (Fig. 1 E) indicating that BK channel activity contributes to basal autophagic flux and supporting the use of BK agonists to enhance this process. Western blotting further corroborated these findings, as starvation reduced the levels of p62 and LC3B-II relative to control cells, consistent with increased autophagic turnover. Notably, inclusion of penitrem A restored these markers to control levels, indicating that BK channel activity contributes to basal starvation-induced autophagic flux (Fig. 1F).

**Fig. 1:**
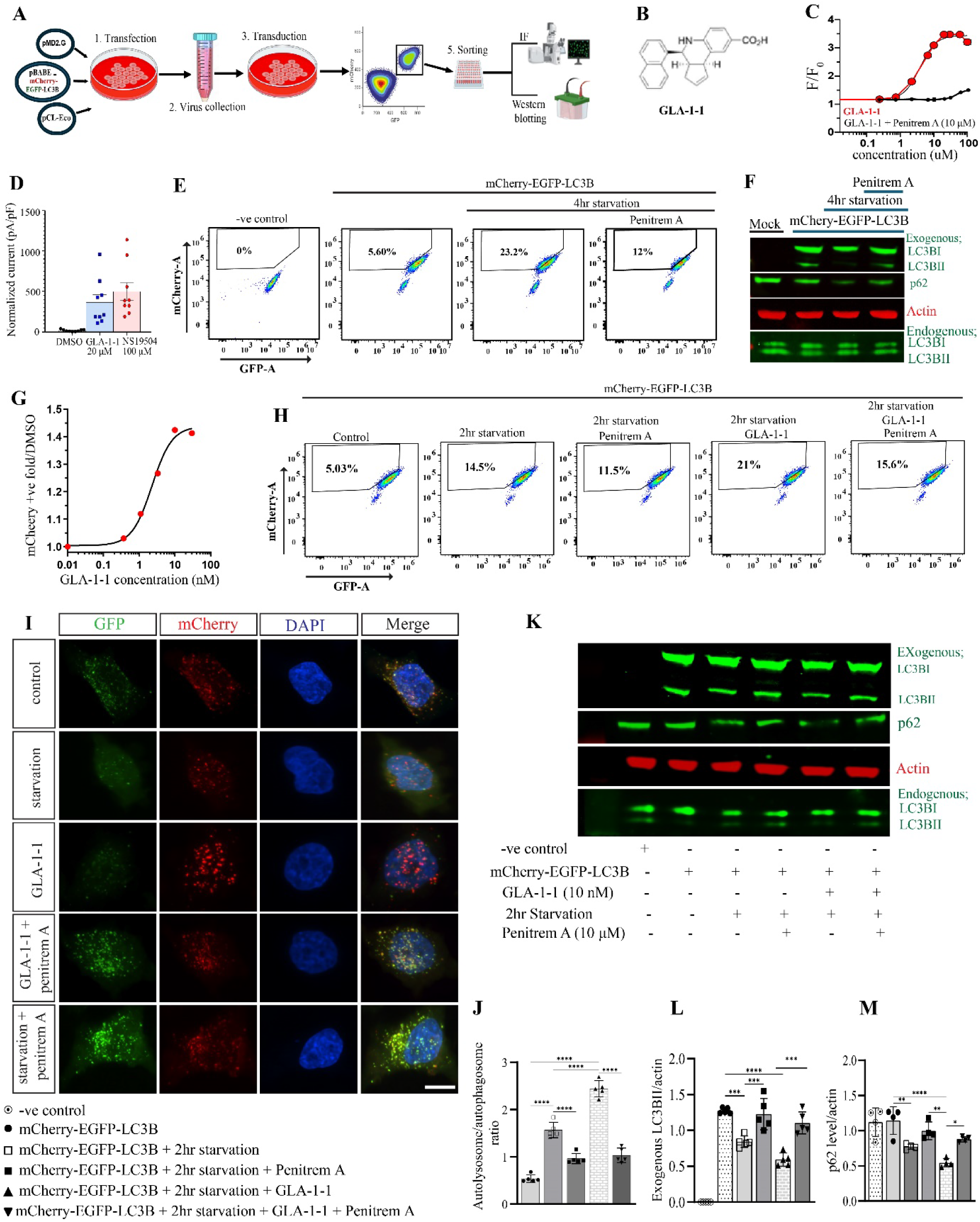
BK agonist GLA-1-1 induces autophagy flux by promoting autophagosome-lysosome fusion. (A) Schemetic representation depicting workflow of reporter cell line generation and autophagy flux induction. (B) Concentration-dependent titration of GLA-1-1 in the FLIPR BK channel agonist assay. Data points represent normalized fluorescence emission of the membrane potential-sensitive dye at different GLA-1-1 concentrations (n = 6 biological replicates and n = 2 technical replicates). The inset shows the structure of GLA-1-1. (C) Effect of BK channel inhibitor penitrem A on GLA-1-1 mediated BK channel activity (n=6 biological replicates and n=2 technical replicates). Data points represent normalized fluorescence emission of the membrane potential-sensitive dye at different GLA-1-1 concentrations in the presence of 50 nM penitrem A. (D) Patch-clamp assay demonstrating GLA-1-1-mediated BK channel activation. Electrophysiological analysis showing normalized increases in BK currents following application of test compounds or DMSO, with individual recording data points shown. The BK agonist NS19504 was included as a positive control and tested at a higher concentration. (E) FACS results suggesting positive role of BK channels in autophagy flux (n=5 biological replicates and n=2 technical replicates). (F) Western blot showing expression of LC3B and p62 in reporter cells in presence and absence of penitrem A. 4-hour serum starvation degrades p62 and LC3BII levels which was significantly inhibited by penitrem A inclusion in starving media (n=3). (G) FACS based titration graph of GLA-1-1 showing autophagy flux from 0.3 nM to 30mM concentration. Maximum flux was observed at 10nM concentration (n=5 biological replicates and n=2 technical replicates). (H) FACS results showing that GLA-1-1 treatment enhances mCherry/GFP ratio show and is sensitive to BK channel inhibitor penitrem A (n=5 biological replicates and n=2 technical replicates). (I) Confocal microscopy results showing GLA1-1 increases in mCherry puncta/ GFP puncta ratio. (J) Quantification of mCherry/GFP ratio of 10 cells per experiment (n=5). (K) Representative western blot depicting effect of GLA-1-1 on autophagy markers p62 and LC3B. (L-M) Relative fold expression of p62 and LC3B (n=5). Statistical analysis was done using one-way ANOVA with post hoc Tukey test for J, L and M. *\*\*\*\*P < 0.0001*.

We next examined the concentration-dependent effect of GLA-1-1 on autophagosome-lysosome fusion and autophagic flux. Reporter cells were treated with increasing concentrations of GLA-1-1 (0.3-30 nM), and flux was quantified by FACS analysis. GLA-1-1 produced a concentration-dependent increase in autophagic flux that peaked at 10 nM (Fig. 1G). Based on this result, 10 nM GLA-1-1 was used to determine whether the observed effect was BK channel-dependent. Serum-starved reporter cells were treated with GLA-1-1 in the presence or absence of the BK channel inhibitor penitrem A. FACS analysis showed a marked increase in the mCherry/GFP fluorescence ratio in cells treated with 10 nM GLA-1-1 compared with control and serum-starved cells. The proportion of cells exhibiting an elevated mCherry/GFP ratio increased from 14.5% under starvation-induced basal autophagy to 21% following GLA-1-1 treatment, representing approximately a 1.5-fold increase in the number of cells with enhanced autophagic flux. Importantly, inclusion of penitrem A significantly attenuated this increase, indicating that the GLA-1-1-induced enhancement of autophagic flux is BK channel-dependent (Fig. 1H). Consistently, confocal microscopy further confirmed these findings. Serum starvation increased the number of red puncta (autolysosomes) relative to baseline, and this effect was further enhanced by GLA-1-1 treatment, consistent with increased fusion and autophagic flux. Importantly, inclusion of penitrem A reduced the number of red puncta toward levels observed with starvation alone (Fig. 2I, J). We next assessed the expression of canonical autophagy markers, including p62/SQSTM1 and LC3B-II. Accumulation of LC3B-II and p62 reflects autophagosome buildup, whereas effective autophagic flux leads to their degradation in lysosomes. Consistent with enhanced autophagic turnover, starvation decreased LC3B-II and p62 levels relative to control cells, an effect that was further enhanced by GLA-1-1 treatment. In contrast, co-treatment with the BK channel inhibitor penitrem A partially reversed this effect, restoring LC3B-II and p62 levels toward those observed with starvation alone, consistent with inhibition of autophagosome-lysosome fusion (Fig. 2K-M). Because BK channel activation in the FLIPR assay occurred at micromolar concentrations, while enhancement of autophagic flux was observed in the nanomolar range, we sought to clarify this difference. Using LC-MS/MS, we measured intracellular concentrations of GLA-1-1 in reporter cells following treatment with 10 nM compound. Strikingly, intracellular GLA-1-1 accumulated to approximately 2 µM, consistent with concentrations required for effective BK channel activation and suggesting efficient cellular uptake and accumulation of GLA-1-1 (Table S1). Together, these findings indicate that GLA-1-1 acts as a potent BK channel agonist that enhances autophagic flux by promoting autophagosome-lysosome fusion.

**Fig. 2:**
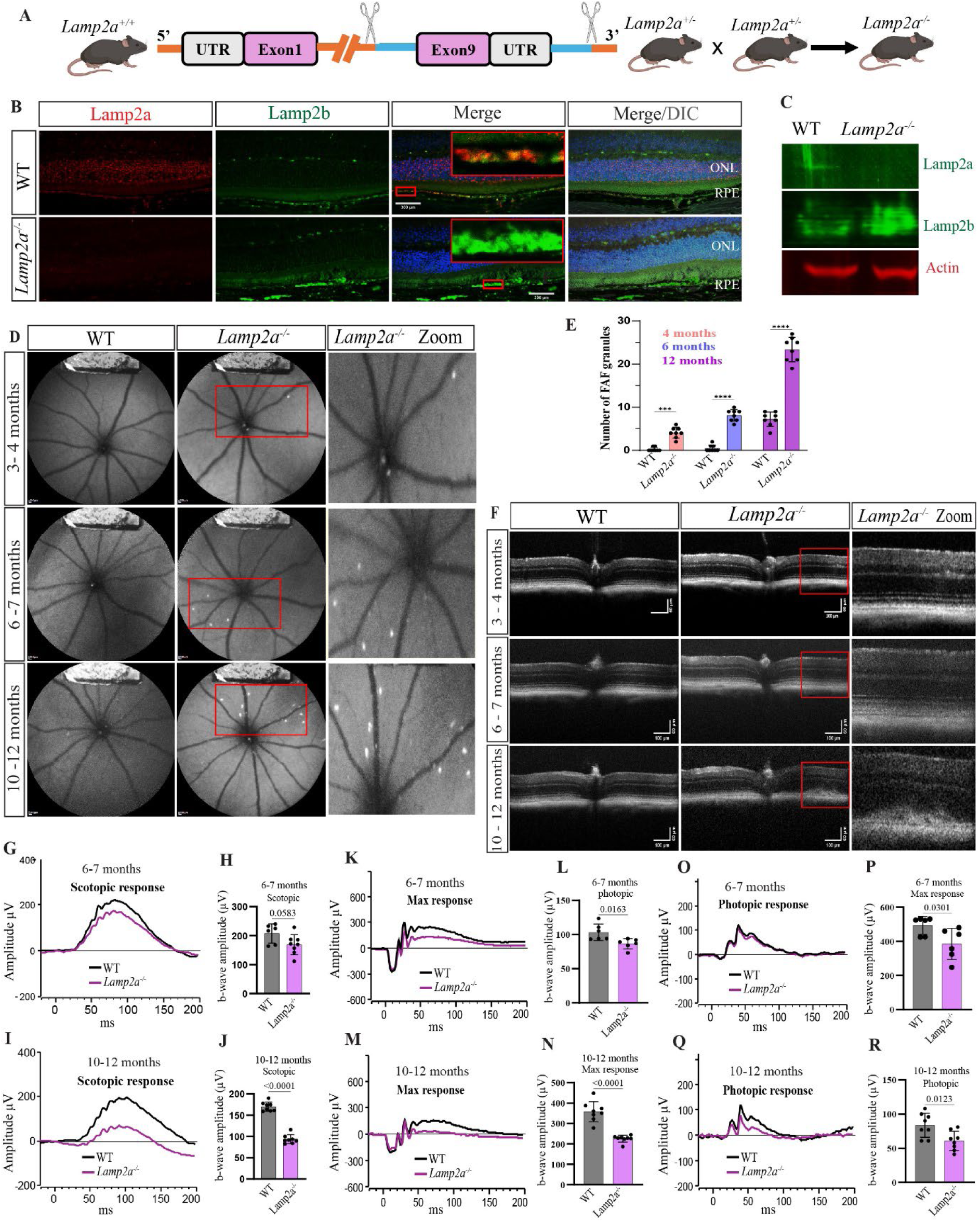
Lamp2a loss shows AMD like age dependent fundus, retinal and visual abnormalities. (A) Cartoon representation of *Lamp2a* knockout mouse generation. (B) Confocal microscopy imaging validating deletion of Lamp2a. Red staining reflects Lamp2a, green staining shows Lamp2b. Scale bar =300 µm (C) Immunoblot of RPE/choroid lysate showing loss of Lamp2a expression in *Lamp2a^-/-^* mice compared to wild type (WT) age matched mice. (D) FAF images showing high-intensity autofluorescence spots in WT and *Lamp2a^-/-^* mice. Lamp2a deficiency results in an age-dependent increase in high-intensity autofluorescence spots compared with WT mice. (E) Quantification of high-intensity autofluorescence spots (n=8). Statistical analysis was performed using Two-way ANOVA with Dunnett’s multiple comparisons test. *\*\*\*P < 0.001; ****P < 0.0001*. (F) Retinal B-scans of *Lamp2a^-/-^* and WT mice. Significant structural disturbance in retinal layers was seen in 12 months *Lamp2a^-/-^* mice as compared to WT aged mice and *Lamp2a^-/-^* younger mice (n=6). Scale bar =100 µm (width) and 60 = µm (height). Values are expressed as mean ± SD. (G) Representative scotopic ERG response of 6-7-month-old WT and *Lamp2a^-/-^* mice. (H) b-wave quantification indicating decline in scotopic response in *Lamp2a^-/-^* mice relative to age-matched wild type (n=6 mice/group). (I) Representative scotopic response of 10-12-month-old WT and *Lamp2a^-/-^* mice. (J) B-wave quantification shows significant decline in scotopic response in *Lamp2a^-/-^* mice relative to age-matched WT (n=8 mice/group). (K) Representative maximal response of 6-7-month-old WT and *Lamp2a^-/-^* mice. (L) b-wave amplitude quantification shows decreased scotopic response in 6-7-month-old *Lamp2a^-/-^*mice relative to age-matched WT (n=6 mice/group). (M) Representative maximal response of 10-12 months old WT and *Lamp2a^-/-^* mice. (N) b-wave amplitude quantification shows significant decrease in scotopic response in 10-12-month-old *Lamp2a^-/-^*mice relative to age-matched wild type (n=8 mice/group). (O) Representative photopic response of 6-7-month-old WT and *Lamp2a^-/-^* mice. (P) b-wave amplitude quantification shows photopic response changes in 6-7-month-old *Lamp2a^-/-^* mice relative to age-matched WT (n=6 mice/group). (Q) Representative photopic response of 10-12-month-old WT and *Lamp2a^-/-^* mice. (R) b-wave amplitude quantification shows pronounced decrease in photopic response in 10-12-month-old *Lamp2a^-/-^* mice relative to age-matched WT (n=8 mice/group). Statistical differences were analyzed using an unpaired two-tailed student’s t-test. Level of significance was considered at p value < *0.05*.

### 2. Selective CMA deficiency induces progressive, age-dependent AMD-like retinal phenotype in *Lamp2a^-/-^* mice

To determine whether CMA deficiency contributes to the retinal phenotype, we studied *Lamp2a^-/-^* mice. Deletion of the Lamp2a isoform in the knockout strain was confirmed by immunofluorescence and Western blotting. Both methods demonstrated loss of Lamp2a expression, accompanied by the increase in Lamp2b expression (Fig. 2B, 2C). Age-dependent retinal changes associated with Lamp2a loss were examined by fundus autofluorescence (FAF) and OCT imaging at 3-4, 6-7, and 10-12 months of age. FAF images showed progressive and significant increase in high-intensity autofluorescence spots compared to age-matched wild type mice (Fig. 2D, E), consistent with the reported retinal phenotype in *Lamp2^-/-^* mice in which all three Lamp2 isoforms were ablated.[15] Punctate or focal areas of increased fundus autofluorescence are a recognized feature of human dry AMD.[43] Retinal structural abnormalities associated with CMA deficiency were further characterized by OCT B-scans (Fig. 2F). While 3-4-month-old *Lamp2a^-/-^* mice were indistinguishable from WT controls, 6–7-month-old *Lamp2a^-/-^* mice displayed occasional RPE hyperplasia which progressed markedly by 12 months and elongated to the outer nuclear layer (ONL) (Fig. 2F), a pattern reminiscent of intraretinal migration of dysmorphic RPE cells into the outer retina described in dry AMD.[44] To determine the effect of CMA deficiency on photoreceptor function, rod-, cone-, and maximal ERG responses were recorded in *Lamp2a^-/-^* mice. Rod-mediated (scotopic) visual responses, indicated by b-wave amplitude in dark-adapted eyes, showed an age-dependent decline in *Lamp2a^-/-^* mice compared with age-matched wild-type controls. While no significant differences were observed in scotopic, max and photopic response in 3–4-month-old mice compared to age matched WT controls (Fig. S2A-F), scotopic sensitivity began to decline at 6 months and was markedly reduced by 12 months in *Lamp2a^-/-^* mice (Fig 2G-J). The maximal scotopic b-wave, representing the mixed rod–cone response at high flash intensities, was also significantly reduced in 12-month-old *Lamp2a^-/-^* mice (Fig. 2K-N). Similarly, photopic responses mediated by cones declined beginning at 6 months, with the decrease becoming more pronounced by 12 months in *Lamp2a^-/-^* mice (Fig. 2O-R). Taken together, these ERG results indicate that CMA deficiency due to Lamp2a loss not only produces structural abnormalities resembling those seen in dry AMD but also leads to comparable visual impairment. Our data indicates that the pathogenic phenotype in *Lamp2a^-/-^* mice progresses with age and is most pronounced at 12 months, making this time point appropriate for testing potential therapeutic interventions.

### 3. BK agonist GLA1-1 reduces autofluorescent deposits and RPE pathology in *Lamp2a^-/-^* CMA-deficient mice

CMA deficiency resulting from LAMP2A abnormalities may be partially compensated by increased macroautophagy. [36, 37] Previous studies have reported compensatory regulation among LAMP2 isoforms, with loss of Lamp2a associated with increased Lamp2b expression, an isoform implicated in macroautophagy.[45, 46] Our confocal staining and immunoblotting results of RPE/choroid samples also confirm increased expression of Lamp2b in *Lamp2a^-/-^*mice in comparison to WT mice (Fig. 2B, C). Given that our in vitro studies showed that GLA-1-1 enhances autophagic flux by promoting autophagosome-lysosome fusion, we investigated whether GLA-1-1 could ameliorate retinal pathology in *Lamp2a^-/-^* mice by enhancing this compensatory autophagy pathway. We treated *Lamp2a^-/-^* mice with GLA-1-1 at the oral dose of 25 mg/kg for 6 months, starting at 6 months of age. FAF imaging showed that GLA-1-1 treatment significantly reduced the accumulation of high-intensity autofluorescent puncta (Fig. 3B, C). OCT retinal B-scans further showed that GLA-1-1 treatment prevented RPE hyperplasia. The extension of dysmorphic RPE cells into the ONL, observed in untreated *Lamp2a^-/-^* mice, was not detected in GLA-1-1–treated *Lamp2a^-/-^* mice (Fig. 3D). Confocal autofluorescence microscopy (488 nm excitation), which detects lipofuscin-associated bisretinoids, showed increased autofluorescent granules in retinal sections of *Lamp2a^-/-^* mice localized to the RPE and sub-RPE regions compared with WT mice (Fig. 3E). Consistent with therapeutic benefit, GLA-1-1–treated *Lamp2a^-/-^*mice showed a significant reduction in the accumulation of autofluorescent granules in the RPE regions (Fig. 3E-G). We also confirmed the oral bioavailability of GLA-1-1 by measuring its serum concentrations using LC–MS/MS following administration via chow. Micromolar concentrations were detected in serum, indicating sufficient systemic exposure to mediate the observed rescuing effects (Table S3). These results demonstrate that important features of the dry AMD phenotype recapitulated in the *Lamp2a^⁻/⁻^* model can be significantly attenuated by pharmacological activation of BK channels with GLA-1-1.

**Fig. 3:**
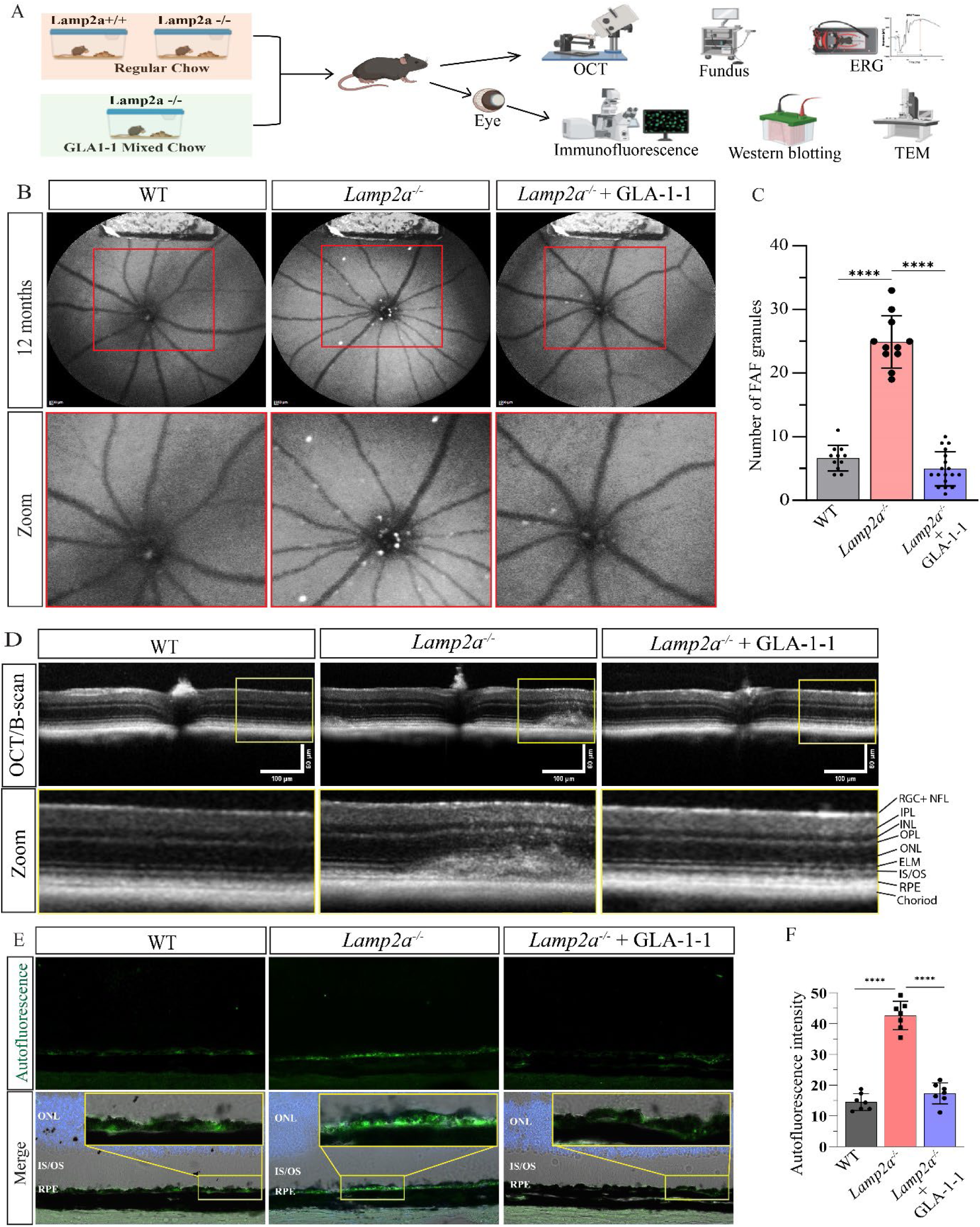
GLA-1-1 reduces autofluorescent deposits and RPE abnormalities in *Lamp2a^-/-^* mice as detected by FAF, OCT, and confocal microscopy. (A) Schematic overview of the experimental design and analytical approaches used in this study. (B) FAF images showing that Lamp2a loss leads to accumulation of high-intensity autofluorescence spots, which are significantly reduced by GLA-1-1 treatment. (C) Quantification of high-intensity autofluorescence spots (WT (n=11), *Lamp2a^-/-^*mice (n=11), *Lamp2a^-/-^* + GLA-1-1 mice (n=18)). (D) OCT B-scans of WT, *Lamp2a^-/-^* untreated, and *Lamp2a^-/-^*-GLA-1-1-treated mouse eyes. GLA-1-1 treatment prevented RPE hyperplasia and extension of dysmorphic RPE cells into the ONL. (E) Autofluorescence images of retinal sections showing accumulation autofluorescent granules in *Lamp2a^-/-^*as compared to WT mice. GLA-1-1 treatment significantly reduced the accumulation of the granules in RPE of *Lamp2a^-/-^* mice. The images were captured with confocal microscope using the excitation wavelength of 488 nm and emission wavelengths of 500–600 nm. (F) Quantification of fluorescence intensity of autofluorescence granules (n=7). Statistical analysis was performed using one-way ANOVA with post hoc Tukey test for C and F ***P < 0.001; ****P < 0.0001. Values are expressed as mean ± SD.

### 4. GLA1-1 ameliorates RPE degeneration and visual impairment in *Lamp2a^-/-^* mice

RPE dysfunction and degeneration are central to dry AMD progression and culminate in geographic atrophy, the hallmark of late-stage disease, characterized by RPE cell loss and associated degeneration and thinning of the overlying photoreceptor layers [47–49]. To characterize anatomical changes in retinal architecture associated with CMA deficiency due to *Lamp2a* loss, we performed histological analysis of *Lamp2a^⁻/⁻^* mouse retinas. We observed substantial enlargement of the IS/OS region in *Lamp2a^-/-^* mice (Fig. 4A), which may reflect accumulation of photoreceptor outer segments (POS) in the subretinal space due to impaired RPE phagocytosis, an effect previously described in the global Lamp2 knockout model [15]. GLA-1-1 treatment prevented the enlargement of the IS/OS region observed in *Lamp2a^-/-^* mice (Fig 4A, B). Loss of Lamp2a resulted in significant thinning of the INL layer relative to WT mice, an effect that was prevented by GLA-1-1 treatment (Fig. 4C). Likewise, the thickness of the RPE, ONL and RGC layers was markedly reduced in *Lamp2a^-/-^* mice compared with WT mice (Fig. 4D; Fig. S3A-B). GLA-1-1 treatment substantially preserved the thickness of both layers (Fig. 4D; Fig. S3A-B).

**Fig. 4:**
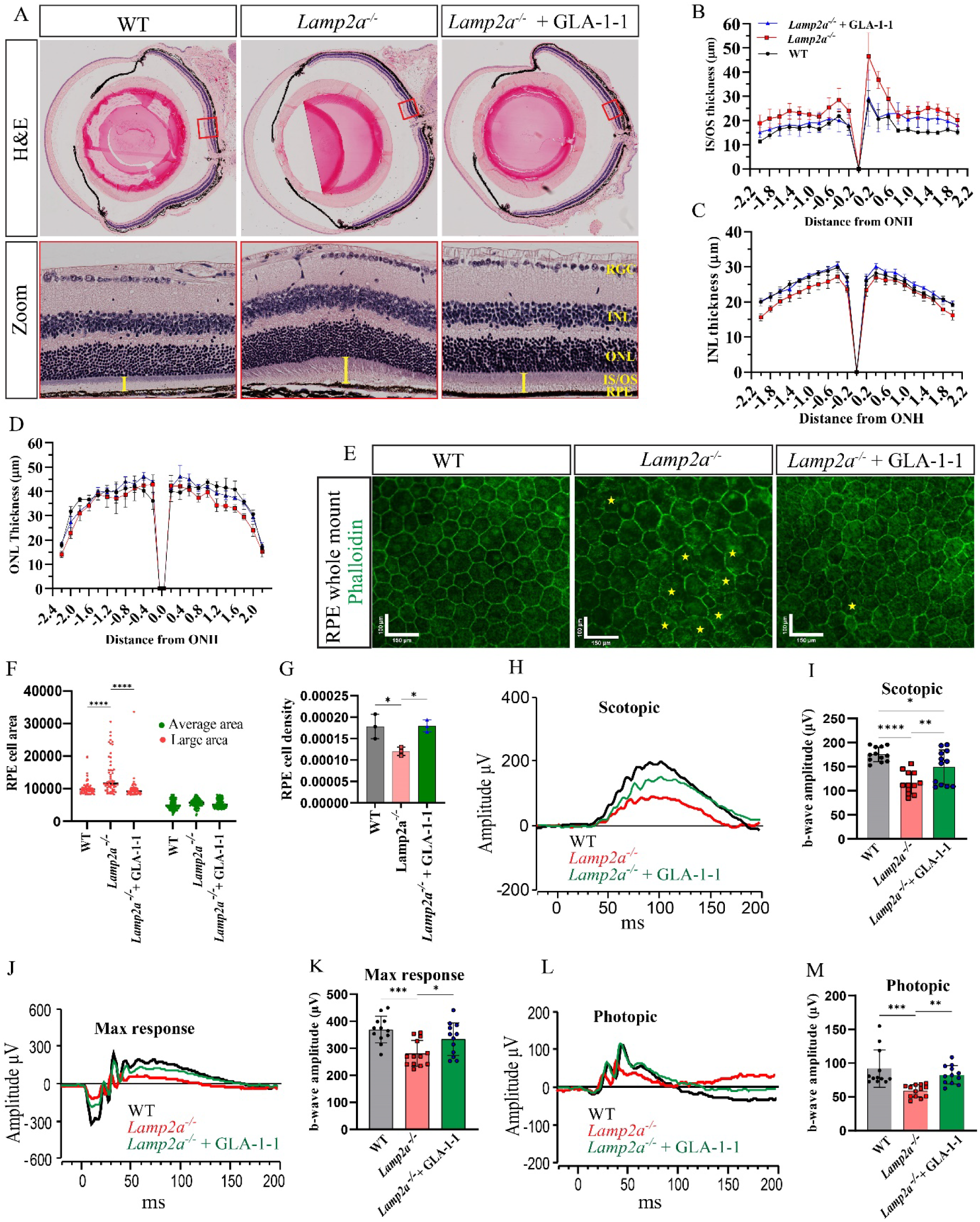
GLA-1-1 preserves retinal structure, RPE morphology, and visual function in *Lamp2a^-/-^* mice. (A) H&E-stained retinal sections showing enlargement of the IS/OS region in *Lamp2a^-/-^* mice. GLA-1-1 treatment prevented this enlargement in *Lamp2a^-/-^* mice. (B) Radial plot showing IS/OS layer thickness measured at 200 µm intervals from the optic nerve head. Thickness was increased in *Lamp2a^-/-^* mice compared with WT mice, and GLA-1-1 treatment reversed this increase. (WT, n=5, *Lamp2a^-/-^* n=6, *Lamp2a^-/-^* + GLA-1-1 n=5). (C-D) Thickness analysis of the INL and ONL layers. Lamp2a loss resulted in a significant decrease in the thickness of both layers, which was markedly attenuated by GLA-1-1 treatment. (WT, n=5, *Lamp2a^-/-^* n=6, *Lamp2a^-/-^*+ GLA-1-1 n=5). (E) Representative phalloidin staining of RPE whole mounts showing RPE cell morphology in WT, *Lamp2a^-/-^*, and GLA-1-1–treated *Lamp2a^-/-^* mice. (F-G) RPE cell area and density quantification from whole mounts showing significant RPE cell loss and enlarged cell areas in *Lamp2a^-/-^* mice, which were prevented by GLA-1-1 treatment. (n=3). Scale bar =150 µm (width) and 100 = µm (height). Statistical analysis was performed using Two-way ANOVA with Dunnett’s multiple comparisons test. *\*\*\*P < 0.001*. (H, J, L) Representative ERG responses showing scotopic rod pathway responses (H), maximal scotopic mixed rod–cone responses at high flash intensities (J), and photopic cone pathway responses (L). (I, K, M) Quantification of b-wave amplitudes showing significant reductions in rod pathway responses (I), maximal mixed rod–cone responses (K), and cone pathway responses (M) in *Lamp2a^-/-^* mice, which were restored by GLA-1-1 treatment. (WT, n=12, *Lamp2a^-/-^*n=13, *Lamp2a^-/-^* + GLA-1-1 n=12). Statistical analysis was performed using one-way ANOVA with post hoc Tukey test. *P < 0.05, **P < 0.01, ***P < 0.001, ****P < 0.0001. Values are expressed as mean ± SD.

Loss of epithelial polarity and disruption of the RPE monolayer, accompanied by migration of dysmorphic RPE cells into the neurosensory retina, are characteristic features of dry AMD progression.[50, 51] To assess these changes in CMA-deficient *Lamp2a^-/-^* mice, we performed whole-mount phalloidin staining of the RPE. In WT mice, the RPE displayed a highly organized morphology with regularly arranged hexagonal cells. In contrast, the RPE in *Lamp2a^-/-^* mice appeared dysmorphic, characterized by enlarged cell areas, loss of clear cell boundaries, and reduced cell density (Fig. 4E-G), features commonly observed in AMD. [52] GLA-1-1 treatment of *Lamp2a^-/-^* mice reduced RPE cell loss and decreased the proportion of cells with enlarged areas (Fig. 4E-G). In addition, we detected occasional migration of RPE cells toward the IS/OS region in *Lamp2a^-/-^* mice, which was not observed following GLA-1-1 treatment (Fig. S3C).

We next examined whether these structural abnormalities and their restoration by GLA-1-1 were reflected in retinal function in CMA-deficient mice using ERG. Scotopic ERG responses reflecting rod pathway function, indicated by b-wave amplitude, declined in *Lamp2a^-/-^* mice compared with WT mice and were restored by GLA-1-1 treatment (Fig 4H, I). The maximal scotopic b-wave, representing the mixed rod–cone response at high flash intensities, was also reduced in *Lamp2a^-/-^*mice and was restored by GLA-1-1 treatment (Fig 4J, K). Similarly, photopic ERG responses reflecting cone pathway function were significantly reduced in *Lamp2a^-/-^* mice compared with WT mice and were restored by GLA-1-1 treatment (Fig 4L, M). Collectively, these results suggest that loss of Lamp2a-mediated CMA in mice leads to structural and functional abnormalities reminiscent of features observed in dry AMD, which can be ameliorated by the BK channel agonist GLA-1-1.

### 5. CMA deficiency in *Lamp2a^-/-^* mice promotes accumulation of drusen-associated proteins and extracellular matrix alterations that are attenuated by GLA-1-1

Drusen, extracellular deposits that accumulate between the RPE and Bruch’s membrane and represent a hallmark lesion of dry AMD, are composed of lipids, proteins, and extracellular matrix components. [53–55] Proteins such as ApoE and the lipid droplet protein perilipin-2 (PLIN2), which are involved in lipid transport and storage, have been identified as components of drusen. [56, 57]. In this context, we examined the expression of these proteins using immunofluorescence and immunoblotting. Immunofluorescence analysis revealed increased ApoE accumulation in the sub-RPE region of *Lamp2a^-/-^* mice, which was not observed in *Lamp2a^-/-^*mice treated with GLA-1-1 (Fig. 5A, B). ApoE staining was also detected in subretinal regions above the RPE, a location corresponding to that of reticular pseudodrusen in human dry AMD (Fig. S4A). Likewise, PLIN2 levels were significantly increased in the sub-RPE region of *Lamp2a^-/-^*mice compared with WT mice, consistent with enhanced lipid accumulation. In contrast, GLA-1-1–treated *Lamp2a^-/-^* mice displayed markedly reduced PLIN2 expression (Fig. 5A, C). Previous studies have reported significant alterations in extracellular matrix proteins in dry AMD, particularly within Bruch’s membrane and drusen-associated deposits.[55, 58, 59] We next examined the distribution of extracellular matrix proteins associated with drusen, including vitronectin and clusterin, using confocal immunofluorescence. Both proteins showed marked accumulation in the sub-RPE region of *Lamp2a^-/-^* mice compared with WT mice. This accumulation was significantly reduced in *Lamp2a^-/-^* mice treated with GLA-1-1. (Fig. 5D, F, G). In contrast, immunoreactivity of certain extracellular matrix proteins, including fibronectin and laminin, was reduced in *Lamp2a^-/-^* mice compared with WT mice, and this reduction was restored by GLA-1-1 treatment (Fig. S4B). We further assessed the protein levels of clusterin, vitronectin, PLIN2, and ApoE by immunoblotting of RPE/choroid lysates. The results were consistent with the immunostaining data, showing elevated levels of these proteins in *Lamp2a^-/-^* mice relative to WT mice. Notably, GLA-1-1 treatment reduced the levels of these proteins in *Lamp2a^-/-^* mice (Fig. 5 H-L). In addition, we examined the localization of matrix metalloproteinase-2 (MMP2) by immunofluorescence. In WT mice, MMP2 immunoreactivity was predominantly detected on the basolateral side of the RPE (sub-RPE region). In *Lamp2a^-/-^* mice, overall MMP2 staining was increased and was observed not only in the sub-RPE region but also on the apical side of the RPE, above the RPE layer. Treatment with GLA-1-1 reduced the overall MMP2 staining and eliminated the apical signal while preserving the basolateral (sub-RPE) localization (Fig. 5D, E). Taken together, these findings indicate that CMA deficiency in *Lamp2a^-/-^* mice leads to accumulation of proteins associated with human drusen together with alterations in lipid handling and extracellular matrix organization, changes that are substantially attenuated by BK channel activation with GLA-1-1.

**Fig. 5:**
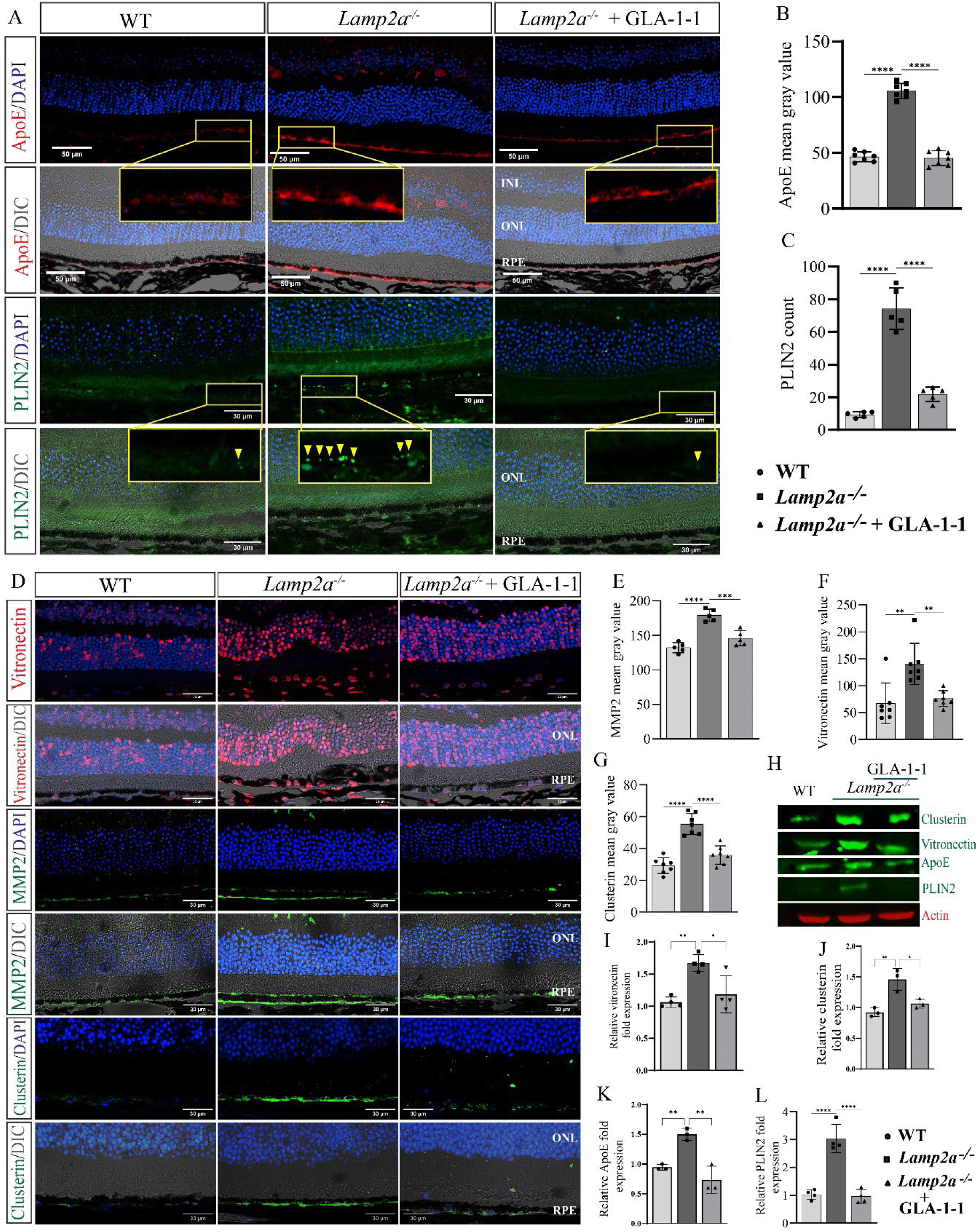
Lamp2a deficiency in mice promotes accumulation of drusen-associated proteins and extracellular matrix alterations that are attenuated by GLA-1-1. (A) Representative confocal immunofluorescence images showing ApoE and PLIN2 staining (green). Scale bar = 30 µm. *Lamp2a^-/-^* mice showed increased staining of both proteins compared with WT mice, which was markedly reduced by GLA-1-1 treatment. (B) Bar graph showing quantification of ApoE fluorescence intensity (mean gray value) (n = 7 mice). (C) Bar graph showing quantification of PLIN2 immunofluorescence signal (n = 5 mice). (D) Representative confocal immunofluorescence images showing staining for vitronectin, MMP2, and clusterin. Scale bar = 30 µm. Compared with WT mice, *Lamp2a^-/-^* mice showed markedly increased immunoreactivity for these proteins, which was significantly reduced by GLA-1-1 treatment. (E) Bar graph showing quantification of MMP2 fluorescence intensity (mean gray value) (n=5 mice). (F) Bar graph showing quantification of vitronectin fluorescence intensity (mean gray value) (n=7 mice). (G) Bar graph showing quantification of clusterin fluorescence intensity (mean gray value) (n=7 mice). (H) Representative immunoblots showing protein levels of ApoE, clusterin, vitronectin, and PLIN2 in RPE/choroid lysates. Bar graphs showing immunoblot quantification of protein levels of vitronectin (I), clusterin (J), ApoE (K), and PLIN2 (L) in RPE/choroid lysates (n = 3 mice per group). Statistical analysis was performed using one-way ANOVA with Tukey’s post hoc test. *P < 0.05, **P < 0.01, ***P < 0.001, ****P < 0.0001. Values are expressed as mean ± SD.

### 6. Lamp2a deficiency promotes RPE lipid accumulation and Bruch’s membrane thickening that are attenuated by GLA-1-1

Dry AMD progression is characterized by accumulation of lipid-rich deposits at the RPE–Bruch’s membrane interface, including soft drusen and basal linear deposits (BLinD), a diffuse lipid-rich material located within the inner collagenous layer of Bruch’s membrane. [53, 60, 61] Given the strong association between these lipid deposits and disease progression, we examined whether *Lamp2a^-/-^* mice develop a similar lipid accumulation phenotype and whether it can be improved by GLA-1-1 treatment. Consistent with this possibility, *Lamp2a^-/-^*mice showed increased accumulation of the lipid-associated proteins ApoE and PLIN2 (Fig. 5A-C), prompting us to further examine lipid accumulation in the RPE and potential alterations in Bruch’s membrane structure. Histochemical analysis revealed marked lipid accumulation in the RPE and sub-RPE regions of *Lamp2a^-/-^* mice. Nile Red and Oil Red O staining demonstrated substantial accumulation of neutral lipids in the RPE and sub-RPE regions of *Lamp2a^-/-^* mice compared with WT mice (Fig. 6A–D). In addition, filipin staining confirmed accumulation of unesterified cholesterol in the sub-RPE region (Fig. S4C, D). In contrast, GLA-1-1–treated *Lamp2a^⁻/⁻^* mice showed a marked reduction in lipid accumulation (Fig. 6A–D). We further examined lipid accumulation in the RPE at the ultrastructural level using transmission electron microscopy (TEM). TEM analysis confirmed extensive lipid accumulation in the RPE of *Lamp2a^-/-^* mice, which was reduced to near WT levels following GLA-1-1 treatment (Fig. 6E, F). Concomitant with lipid accumulation, TEM analysis also demonstrated a significant increase in Bruch’s membrane thickness in *Lamp2a^-/-^* mice, which was attenuated by GLA-1-1 treatment (Fig. 6G, H). The increase in Bruch’s membrane thickness was further confirmed by periodic acid–Schiff (PAS) staining, which showed pronounced thickening in *Lamp2a^-/-^*mice. Notably, GLA-1-1 treatment reduced Bruch’s membrane thickness to levels comparable to WT mice (Fig. 6I, J). Together, these findings demonstrate that CMA deficiency in *Lamp2a^-/-^* mice leads to lipid dysregulation characterized by accumulation of neutral lipids and cholesterol in the RPE and sub-RPE regions, accompanied by Bruch’s membrane thickening, changes reminiscent of lipid-rich deposits observed in dry AMD, which are markedly attenuated by GLA-1-1 treatment.

**Fig. 6:**
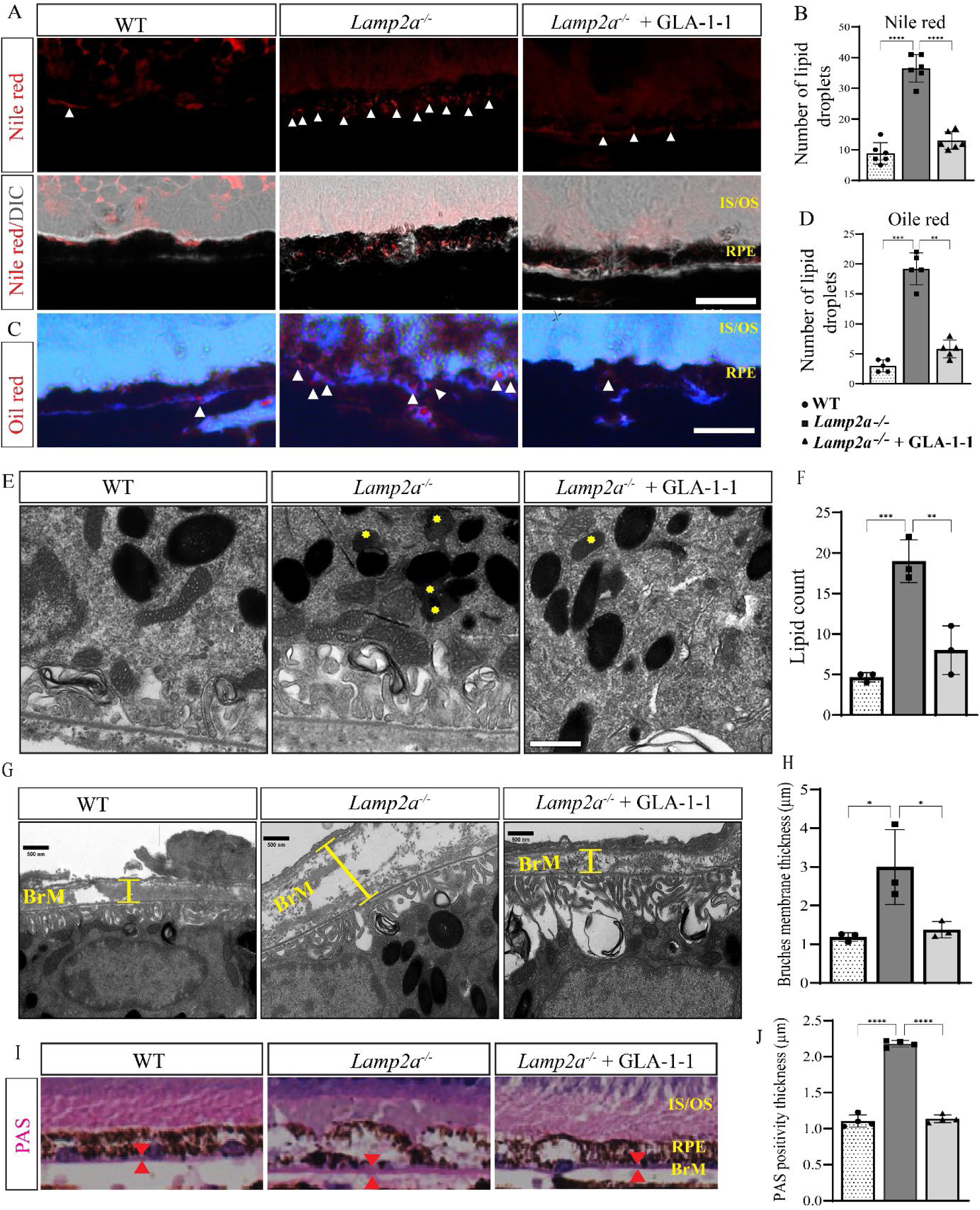
GLA-1-1 attenuates Lamp2a deficiency-associated lipid accumulation in the RPE and reduces Bruch’s membrane thickening. (A) Representative Nile Red staining showing neutral lipid droplets in the RPE and sub-RPE regions. Scale bar = 200 µm. (B) Quantification of Nile Red–positive lipid droplets showing increased neutral lipid accumulation in *Lamp2a^-/-^* mice compared with WT mice, which was markedly reduced by GLA-1-1 treatment (n = 6 mice). (C) Representative Oil Red O staining showing accumulation of neutral lipids in the RPE and sub-RPE regions. Scale bar = 200 µm. (D) Quantification of Oil Red O-positive lipid droplets showing increased lipid accumulation in *Lamp2a^-/-^* mice, which was significantly reduced by GLA-1-1 treatment (n = 5 mice). Transmission electron microscopy (TEM) images showing intracellular lipid droplets (yellow asterisks) in the RPE of Lamp2a⁻/⁻ mice. Lipid accumulation was reduced following GLA-1-1 treatment. Scale bar = 1 µm. (F) Quantification of lipid droplets observed in TEM images (n = 3 mice). (G) Representative TEM images showing Bruch’s membrane (BrM). Scale bar = 500 nm. (H) Quantification of Bruch’s membrane thickness showing significant thickening in *Lamp2a^-/-^* mice compared with WT mice, which was attenuated by GLA-1-1 treatment (n = 3 mice). Thickness was measured every 2 µm in three different fields per mouse. (I) Periodic acid–Schiff (PAS) staining of retinal sections highlighting Bruch’s membrane thickening in *Lamp2a^-/-^*mice. (J) Quantification of PAS-positive Bruch’s membrane thickness showing significant thickening in *Lamp2a^-/-^* mice, which was markedly reduced by GLA-1-1 treatment (n = 4 mice). Thickness was measured every 200 µm. Statistical analysis was performed using one-way ANOVA with Tukey’s post hoc test. *P < 0.05, **P < 0.01, ***P < 0.001, ****P < 0.0001. Values are expressed as mean ± SD.

### 7. CMA deficiency in *Lamp2a^⁻/⁻^* mice disrupts lysosome-dependent POS phagocytosis and autophagic flux and is rescued by GLA-1-1

Lysosomal fusion, which can be facilitated by BK channel activation, represents a common convergence point for both autophagy and photoreceptor outer segment (POS) phagocytosis, as lysosomes serve as the terminal degradative compartment for these pathways. We extended the TEM analysis to assess the effects of Lamp2a ablation and GLA-1-1 treatment on RPE phagocytosis. TEM analysis revealed a marked increase in the number of phagosomes in CMA-deficient mice together with significant morphological abnormalities in these structures (Fig. 7A). Quantitative analysis confirmed a significant increase in abnormal phagosomes in *Lamp2a^-/-^* mice compared with WT mice (Fig. 7B). In contrast, *Lamp2a^-/-^* mice treated with GLA-1-1 showed a reduction in abnormal phagosomes and restoration of phagosome morphology toward WT levels (Fig. 7A, B). Because lysosomal dysfunction can also affect mitochondrial turnover, we examined mitochondrial morphology in the same TEM sections. *Lamp2a^-/-^* mice exhibited increased numbers of mitochondria localized at the basal side of the RPE with elongated tubular morphology (Fig. 7A). Quantification confirmed a significant increase in tubular mitochondria in *Lamp2a^-/-^*mice compared with WT mice (Fig. 7C). These mitochondrial abnormalities were markedly reduced in *Lamp2a^-/-^* mice treated with GLA-1-1, consistent with restoration of mitochondrial homeostasis (Fig. 7A, C). These ultrastructural abnormalities, consistent with impairment of lysosome-dependent degradative pathways, prompted us to examine markers of autophagic activity in *Lamp2a^-/-^* mice. We analyzed the autophagy markers LC3B and SQSTM1/p62 by immunostaining and immunoblotting, building on our earlier in vitro observations that BK channel activation promotes autophagosome–lysosome fusion. Immunostaining revealed accumulation of total LC3B in *Lamp2a^-/-^* mice relative to WT mice; SQSTM1/p62 levels were likewise markedly increased (Fig. 7D). The concurrent elevation of LC3B and p62 is consistent with impaired autophagic flux, suggesting that although compensatory macroautophagy may be induced in response to CMA deficiency, degradation through the lysosomal pathway is inefficient. Notably, SQSTM1/p62 accumulation was concentrated in the IS/OS region, where we previously observed reticular pseudodrusen–like autofluorescence, thickening of the IS/OS region, and ApoE accumulation on the apical side of the RPE, consistent with localized impairment in regions exhibiting RPE dysfunction. GLA-1-1 treatment reduced SQSTM1/p62 and total LC3B levels in *Lamp2a^-/-^* mice toward WT levels. (Fig. 7D). Impaired autophagic flux in CMA-deficient mice was further supported by immunoblot analysis of RPE/choroid lysates. Compared with WT mice, *Lamp2a^-/-^*mice showed increased levels of LC3B-II and SQSTM1/p62, consistent with accumulation of autophagic substrates. Treatment with GLA-1-1 significantly reduced p62 levels in *Lamp2a^-/-^*mice. In addition, both LC3B-I and LC3B-II levels were reduced following GLA-1-1 treatment, indicating restoration of autophagic turnover and consistent with our earlier in vitro observations that BK channel activation promotes autophagosome–lysosome fusion (Fig. 7E–G). Consistent with impaired autophagic clearance and accumulation of cellular debris, we detected notable accumulation and activation (amoeboid-like morphology) of Iba1⁺ cells in RPE whole mounts of *Lamp2a^-/-^* mice (Fig. 7H, I), along with their translocation to the IS/OS region in retinal sections (Fig. S5A). In dry AMD, accumulation of Iba1⁺ microglia/macrophages is frequently observed in the subretinal space, where these cells associate with lipid-rich deposits and drusen-like material. [62, 63]. The enrichment of Iba1⁺ cells in the IS/OS region, together with bulging of the IS/OS segment, accumulation of autofluorescent granules, and apical ApoE deposition in the subretinal space, is consistent with the presence of reticular pseudodrusen-like subretinal deposits associated with CMA dysfunction. Notably, GLA-1-1 treatment not only reduced Iba1^+^ cells mice (Fig. 7H, I) but also prevented their subretinal migration (Fig. S5A). Overall, these findings show that CMA deficiency in *Lamp2a^-/-^* mice disrupts lysosome-dependent POS phagocytosis and autophagic flux in the RPE and promotes subretinal inflammatory responses, phenotypes reminiscent of pathogenic processes associated with dry AMD that are markedly attenuated by BK channel activation with GLA-1-1.

**Fig. 7.**
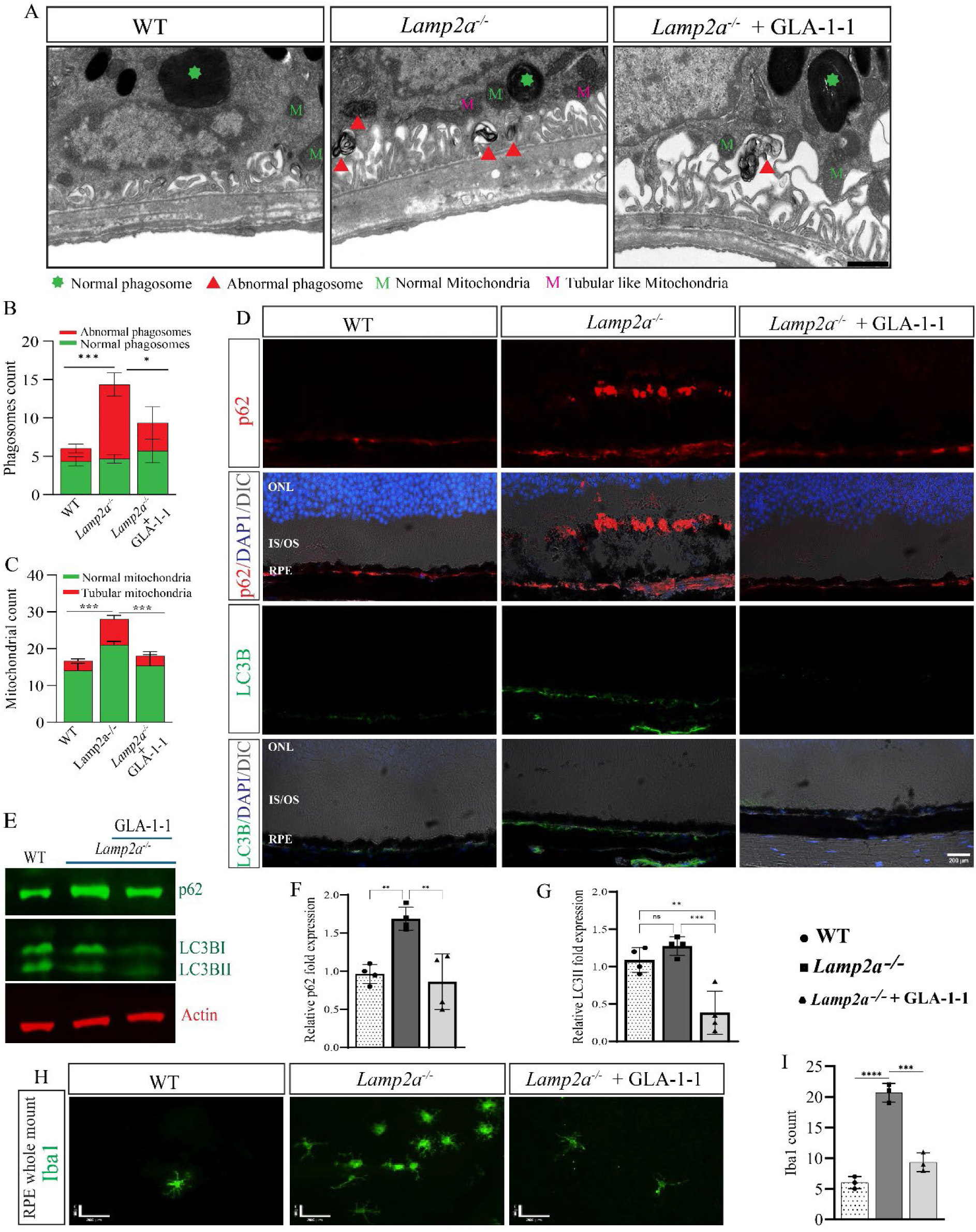
BK channel agonist GLA-1-1 restores lysosome-dependent degradative abnormalities in CMA-deficient *Lamp2a^-/-^* mice. (A) Representative TEM images of the RPE showing increased abnormal phagosomes and elongated tubular mitochondria in Lamp2a⁻/⁻ mice compared with WT mice. These ultrastructural abnormalities were reduced by GLA-1-1 treatment. Green asterisks indicate normal phagosomes, red arrowheads indicate abnormal phagosomes, green letters indicate normal mitochondria, and magenta letters indicate tubular mitochondria. Scale bar = 1 µm. (B) Quantification of normal and abnormal phagosomes (n = 3 mice). (C) Quantification of normal and tubular mitochondria (n = 3 mice). Phagosomes and mitochondria were quantified in three different fields per mouse, and mean values were plotted. (D) Representative confocal immunofluorescence images showing p62 (red) and LC3B (green) staining in retinal sections. *Lamp2a^-/-^* mice showed increased p62 and LC3B immunoreactivity compared with WT mice, which was reduced by GLA-1-1 treatment. DAPI, blue. Scale bar = 200 µm. (E) Representative immunoblots of p62, LC3B-I, LC3B-II, and actin in RPE/choroid lysates. (F) Immunoblot quantification of p62 levels (n = 3 mice). *Lamp2a^-/-^*mice showed increased p62 levels, which were reduced by GLA-1-1 treatment. (G) Immunoblot quantification of LC3B levels (n = 3 mice). *Lamp2a^-/-^* mice showed increased LC3B levels, and GLA-1-1 treatment reduced both LC3B-I and LC3B-II. (H) Representative RPE whole-mount immunostaining showing accumulation of Iba1⁺ cells in *Lamp2a^-/-^* mice, which was reduced by GLA-1-1 treatment. Scale bar = 200 µm (width) and 100 µm (height). (I) Quantification of Iba1⁺ cells in RPE whole mounts (n = 3 mice Statistical analysis was performed using one-way ANOVA with Tukey’s post hoc test. *P < 0.05, **P < 0.01, ***P < 0.001, ****P < 0.0001. Values are expressed as mean ± SD.

## Discussion

Lysosomal dysfunction and impaired autophagy have emerged as important contributors to the pathogenesis of dry age-related macular degeneration (AMD). Previous *in vivo* studies demonstrated that global deletion of *Lamp2*, which eliminates all Lamp2 isoforms, produces age-dependent retinal abnormalities including sub-RPE deposits, Bruch’s membrane thickening, and accumulation of drusen-associated proteins, establishing that disruption of lysosome-dependent degradation in the RPE can drive AMD-like pathology [15]. However, because this model ablates all LAMP2 isoforms, it does not resolve the specific contribution of chaperone-mediated autophagy (CMA), which depends on the LAMP2A isoform. Complementary evidence from human studies implicated CMA in AMD pathogenesis. Cuervo and colleagues reported reduced LAMP2A expression and diminished CMA activity in RPE from AMD patients, accompanied by accumulation of CMA substrate proteins and impaired proteostasis [25]. Despite these advances, it remained unclear whether selective impairment of CMA is sufficient to drive AMD-like pathology *in vivo*. Here we show that loss of Lamp2a in mice produced structural, functional, and molecular changes that resemble key features of human dry AMD. Fundus imaging revealed age-dependent accumulation of autofluorescent puncta in *Lamp2a^-/-^* mice (Fig. 2D, E). OCT and autofluorescence imaging of retinal sections localized these granules predominantly to the sub-RPE region, with additional deposits detected in the subretinal space, resembling the distribution of sub-RPE drusen and reticular pseudodrusen observed in geographic AMD (Fig. 3D–F). Histological analyses further demonstrated pronounced disruption of retinal architecture in *Lamp2a^-/-^*mice, including thinning of the ONL, INL, and RGC layers (Fig. 4A, C, D; Fig. S3B). In addition, abnormal thickening and bulging of the photoreceptor IS/OS region were observed, extending toward the ONL and INL (Fig. 4B). These regions contained accumulated autofluorescent material and ApoE deposits (Fig. S4A), suggesting the presence of drusen-like material. In addition, dysmorphic RPE cells exhibited anterior displacement toward the photoreceptor IS/OS layer, producing localized bulging at the RPE–photoreceptor interface that extended into the ONL and INL (Fig. S3A, C). Whole-mount analysis of the RPE revealed cell loss and increased cellular area in *Lamp2a^-/-^* mice (Fig. 4E–G), consistent with compensatory hypertrophy of surviving RPE cells following neighboring cell loss. Consistent with the central role of RPE cells in maintaining photoreceptor function and visual cycle homeostasis, these structural abnormalities were accompanied by impaired retinal function, including reduced scotopic and photopic ERG responses and a diminished maximal combined ERG response (Fig. 4H–M). In addition to structural and functional abnormalities, Lamp2a deficiency resulted in accumulation of lipids and drusen-associated proteins in the RPE and sub-RPE regions. ApoE and PLIN2, proteins involved in lipid transport and storage, were increased in *Lamp2a^-/-^* mice relative to WT controls (Fig. 5A–C). Likewise, extracellular matrix proteins associated with human drusen, including vitronectin and clusterin, accumulated in the sub-RPE region (Fig. 5D–G). Histochemical staining and ultrastructural analyses further revealed substantial accumulation of lipid droplets in the RPE and sub-RPE space (Fig. 6A–F; Fig. S4C, D), accompanied by thickening of Bruch’s membrane (Fig. 6G–J). Together, these findings indicate that CMA deficiency disrupts lipid and protein clearance pathways in the RPE, leading to accumulation of sub-RPE deposits reminiscent of those observed in early AMD. Given the central role of lysosome-dependent degradation in both autophagy and photoreceptor outer segment phagocytosis, we examined the underlying mechanisms of these pathological changes. Lamp2a deficiency was associated with increased levels of p62 and LC3B, consistent with impaired autophagic flux (Fig. 7D–G). Ultrastructural analysis revealed accumulation of phagosomes with abnormal morphology in the RPE of *Lamp2a^-/-^* mice (Fig. 7A, B), suggesting defective lysosome-dependent processing of phagocytosed POS material. In parallel, we observed accumulation and migration of Iba1⁺ microglial cells toward the IS/OS region (Fig. 7H; Fig. S4A), consistent with recruitment of inflammatory cells to sites of subretinal debris accumulation. Lamp2a deficiency also affected mitochondrial homeostasis, with increased numbers of elongated tubular mitochondria clustered at the basal side of the RPE (Fig. 7A, C). Similar mitochondrial abnormalities have been reported in AMD patient samples and experimental RPE models and are thought to contribute to oxidative stress and retinal degeneration.[64] Our findings, together with the previous report [25], suggest that CMA dysfunction may represent a significant step in the cascade of cellular processes leading to AMD-like pathology. Because effective and convenient therapies for dry AMD are still lacking, we investigated whether pharmacological modulation of lysosome-dependent degradation could ameliorate these defects. BK channels regulate lysosomal ion homeostasis by facilitating Ca²⁺ release from lysosomes through a K⁺-dependent counter-ion mechanism [28–30]. This lysosomal Ca²⁺ signaling promotes SNARE-mediated autophagosome–lysosome fusion and efficient cargo degradation [31, 33]. Consistent with this mechanism, genetic or pharmacological activation of BK channels enhances lysosomal Ca²⁺ release and rescues defects in lysosomal trafficking and degradation [28, 30]. Based on the premise that CMA deficiency may be partially compensated by activation of other degradative pathways such as macroautophagy [36, 37], we first tested the BK channel activator GLA-1-1 in a HeLa reporter assay of autophagosome-lysosome fusion and subsequently evaluated its efficacy in the *Lamp2a^-/-^* mouse model that recapitulates key features of the dry AMD phenotype. We found that the BK channel agonist GLA-1-1 potently activated BK channels and enhanced autophagic flux in the mCherry-EGFP-LC3B reporter cell line (Fig. 1B–D, G–H). Importantly, GLA-1-1 treatment rescued pathological features observed in *Lamp2a^-/-^* mice, including structural retinal abnormalities, visual dysfunction, lipid accumulation, and mitochondrial defects (Fig. 4–7). Mechanistically, GLA-1-1 enhanced autophagic degradation, as reflected by reduced p62 accumulation and decreased LC3B-II levels (Fig. 7D–G), consistent with improved autophagosome–lysosome fusion and enhanced autophagic degradation.

In summary, our study demonstrates that loss of Lamp2a-dependent CMA disrupts lysosome-dependent degradative pathways in the RPE, leading to retinal pathology reminiscent of dry AMD. Pharmacological activation of BK channels restored autophagic flux and attenuated multiple disease-associated phenotypes. These findings highlight the importance of CMA in maintaining protein homeostasis in the RPE and suggest that enhancing lysosome-dependent clearance through BK channel activation may represent a promising therapeutic strategy for dry AMD.

## Materials and Methods

### Compounds, reagents, plasmids and antibodies

GLA-1-1 was synthesized using the published imino Diels-Alder route, which provides the *endo* cycloadduct exclusively [42, 65]. Its identity and purity (>95%) was confirmed by LC-MS, ^1^H NMR, and HPLC analyses; there was no single impurity over 0.5% identified; the compound was proven to be racemic (no optical rotation observed via polarimeter). Our 1D ^1^H NMR was consistent with the reported spectrum [42]. We also further confirmed the relative stereochemistry of GLA-1-1 via a suite of NMR experiments that included DEPT, COSY, HMBC, HMQC, and NOSEY experiments. The NOSEY 2D spectrum shows NOEs between all three hydrogens located on the contiguous asymmetric centers, providing evidence that they are indeed *cis* to one another. NS1619, NS11021 (4788), MaxiPost (4949) and Penitrem A (4617) were obtained from Tocris Bioscience. Polybrene (107689), 4′,6-diamidino-2-phenylindole (DAPI, D1306) and Bafilomycin A1 (SML1661) were purchased from Sigma. Dulbecco’s modified Eagle medium (DMEM, 10313021), fetal bovine serum (FBS), Ham’s F-12 nutrient mix (11765054), Earle’s balanced salt solution (EBSS, 14-155-063), Dulbecco’s phosphate buffered saline (DPBS), Lipofectamine 300 transfection reagent (L3000015), polyvinylidenedifluoride (PVDF, LC2002) membrane and protease inhibitor cocktail were obtained from Thermo Scientific. Antibodies against LC3B (Sigma, L8918-200UL), actin (Proteintech, 66009-1-Ig), PLIN2 (Proteintech, 15294-1-AP), fibronectin (Invitrogen, PA5-29578), clusterin (R&D systems, AF2747), vitronectin (R&D Systems, MAB38751), and p62 (Abnova, H00008878-M01; CST, 5114), Lamp2a (Abcam, ab125068), Lamp2b (Abcam, ab13524), Iba1 (Fujifilm Wako, 019-19741), Laminin (Sigma, L9393) and ApoE (Abcam, ab183596) were used in this study. IR-conjugated (926-68070) anti-mouse and Alexa-flour conjugated secondary (A21206) anti-rabbit were purchased from Licor and Invitrogen respectively.

### Cell culture

HeLa (CCL-2 ™), Human Kca1.1 (BK)/beta1-CHO (Charles River #CT6122), Ampho phoenix (CRL-3213 ™) cells were obtained from ATCC. HeLa and Ampho cells were grown in DMEM containing 10% FBS and 1% penicillin/streptomycin. CHO cells were grown in Hams F12 media containing 10%FBS, 1% penicillin/streptomycin, 0.25 mg genticin (15750060) and 0.1 mg/ml zeocin (invivoGen #ant-zn-5b). All the cell lines were grown and maintained in 5% CO2 incubator at 37 ^0^C.

### FLIPR assay / Membrane potential assay

CHO cell line stably expressing the pore forming Slo1 and the auxiliary β1 subunits was purchased form Charles River (CT6122). We established a 96-well format FLIPR assay using the Membrane Potential Blue dye from Molecular Devices (R8042) as previously described [42]. Cells were pre-incubated with the blue dye and a test compound in the low-K^+^ buffer; after the fluorescence baseline was recorded for 10 sec (530 nm ex/570 nm em), the high-K^+^ stimulus buffer was added, and fluorescence emission recorded for an additional 90 sec. Fluorescence was recorded on BMG LABTECH NOVOstar (model 0700-101) multifunctional microplate reader. The change in fluorescence emission (F/F_0_) was calculated as a ratio of emission at timepoints following the addition of the stimulus buffer to the averaged baseline emission. The F/F_0_ values for the 75 sec timepoint (plateau level) were used for plotting the titration curves and EC_50_ calculations

### Preparation of tandem pBABE-mCherry–EGFP–LC3B retroviral vector

Human AmphoPhoenix cells were used to generate retroviral particles expressing the tandem mCherry–EGFP–LC3B construct. Briefly, AmphoPhoenix cells (3 × 10⁵ cells per well) were seeded in 6-well plates. At approximately 70% confluence, cells were co-transfected with 0.25 μg of envelope plasmid pMDG (Addgene, 12259), 0.75 μg of packaging plasmid pCL-Eco (Addgene, 12371), and 1 μg of pBABE-mCherry–EGFP–LC3B plasmid using Lipofectamine 3000 according to the manufacturer’s instructions. After 18 h, the medium was replaced with fresh culture medium. Retrovirus-containing supernatants were collected at 24 and 48 h after transfection and filtered through a 0.45 μm syringe filter. Viral titters were determined using the QuickTiter Retrovirus Quantification Kit (Cell Biolabs, VPK-120), and target cells were transduced with virus corresponding to approximately 4 × 10⁴ infectious units per cell.

### Generation of autophagic flux reporter HeLa single-cell clone

HeLa cells were seeded in 6-well plates for generation of a stable mCherry–EGFP–LC3B autophagic flux reporter cell line. At 60–70% confluence, the medium was replaced with 1 ml of fresh growth medium supplemented with 1 ml of retrovirus-containing medium carrying the mCherry–EGFP–LC3B construct. Polybrene (8 μg ml⁻¹) was added to enhance viral transduction. Following overnight incubation, the medium was replaced with fresh growth medium containing puromycin (1 μg ml⁻¹) for selection for 48–72 h. Surviving cells were then sorted by flow cytometry for double-positive mCherry and EGFP expression. Single cells were sorted into 96-well plates to generate clonal populations, and bulk transduced cells were collected separately in 15 ml tubes. Successful transduction and expression of the reporter construct were confirmed by western blotting.

### Flow cytometry

HeLa single-cell clones stably expressing the mCherry–EGFP–LC3B reporter were screened for autophagic flux by flow cytometry, and a clone with moderate reporter expression was selected for subsequent experiments. To assess the effect of GLA-1-1 on autophagic flux, the selected HeLa clone was seeded in 6-well plates and grown to 50–60% confluence. The following day, cells were treated with the indicated concentrations of GLA-1-1 overnight in complete growth medium, followed by incubation in EBSS for 2 h on the day of the assay. Maximal autophagic flux was observed at 10 nM GLA-1-1; therefore, this concentration was used in subsequent experiments. In some experiments, cells were co-treated with bafilomycin A1 or penitrem A during the 2-h EBSS incubation. Following treatment, cells were washed three times with DPBS, trypsinized for 1 minute, and collected by centrifugation at 1,200 rpm for 4 min in 15-ml tubes. Cells were then resuspended in 2 ml DPBS and centrifuged again under the same conditions. Finally, cells were resuspended in FACS buffer (EBSS supplemented with 1% FBS), kept on ice, and stained with DAPI (1 μg ml⁻¹) immediately before flow cytometry analysis.

### Confocal Microscopy

HeLa cells stably expressing mCherry-EGFP-LC3B reporter were seeded at low density on glass coverslips. Cells were treated overnight with GLA-1-1 (10 nM) in complete growth medium. The following day, the medium was removed and cells were washed with PBS, followed by incubation in EBSS for 2 h with either GLA-1-1 alone or in combination with penitrem A. After treatment, cells were washed three times with PBS and fixed in 4% paraformaldehyde for 10–15 min. Fixed cells were washed with PBS and mounted in Vectashield mounting medium containing DAPI. Images were acquired using a Nikon Ti Eclipse confocal microscope equipped with a ×40 oil-immersion objective. mCherry–EGFP–LC3B puncta were quantified using Fiji (ImageJ).

### Animal experiments

All animal procedures were approved by the Institutional Animal Care and Use Committee (IACUC) of Columbia University and were conducted in accordance with the Association for Research in Vision and Ophthalmology (ARVO) Statement for the Use of Animals in Ophthalmic and Vision Research. Adult male Lamp2a knockout mice (*Lamp2a^y/-^*), female heterozygous *Lamp2a^+/-^* and their wild type (WT) littermates (*Lamp2a ^y/+^)* were obtained from Cyagen (Serial Number KOCMP-16784-Lamp2-B6J-VA; strain C57BL/6J-Lamp2^em1Cya^). Mice were bred for at least five generations to expand and generate Lamp2a homozygous *Lamp2a^-/-^* female and hemizygous *Lamp2a ^y/-^*male. The GLA-1-1 chow formulation was designed to deliver an approximate oral dose of 25 mg kg⁻¹ day⁻¹. Mouse genotypes were confirmed by standard PCR using primers listed in Table S2.

### In vivo retinal imaging: fundus autofluorescence (FAF) and OCT

For in vivo retinal imaging, mice were anesthetized by intraperitoneal injection of ketamine (100 mg kg⁻¹) and xylazine (10 mg kg⁻¹), followed by pupillary dilation with topical 1% tropicamide and 2.5% phenylephrine hydrochloride. Fundus autofluorescence (FAF) images were acquired using a confocal scanning laser ophthalmoscope (Spectralis HRA; Heidelberg Engineering, Heidelberg, Germany) equipped with a 55° lens and 488 nm excitation. B-scan images of the retina (1.8 mm radial and rectangular volume scans) were obtained at high resolution using spectral-domain optical coherence tomography (SD-OCT; Bioptigen, Leica Microsystems, Buffalo Grove, IL, USA).

### Electroretinography (ERG)

Electroretinography was performed on mice that had been dark-adapted overnight. Mice were anesthetized by intraperitoneal injection of ketamine (100 mg kg⁻¹) and xylazine (10 mg kg⁻¹), followed by pupillary dilation with topical 1% tropicamide and 2.5% phenylephrine hydrochloride. Body temperature was maintained at 37 °C using a heating pad during the recordings. ERG responses were recorded simultaneously from both eyes. Corneal responses were recorded using gold wire loop electrodes placed on the corneal surface and maintained with a drop of Gonak solution. Scotopic ERG responses were recorded under dark-adapted conditions using flashes of 0.001 cd·s m⁻² to measure rod-driven responses, and 3 cd·s m⁻² flashes to measure the maximal combined rod–cone response (white light, 6500 K). Mice were then light-adapted in a Ganzfeld dome for 10 min before recording photopic responses. Cone-mediated photopic ERG responses were recorded using flashes of 30 cd·s m⁻² (xenon) in the presence of a rod-saturating background illumination of 30 cd m⁻² (white, 6500 K).

### Immunostaining, hematoxylin and eosin (H&E) staining, and lipid staining

Eyes were enucleated from freshly euthanized mice, embedded in OCT compound, and frozen in liquid nitrogen. Cryosections (10 μm) were collected on glass slides and fixed in 4% paraformaldehyde for 10 min at room temperature. Sections were washed with PBS for 10 min and blocked with 5% BSA in PBST (0.3% Triton X-100) for 1 h at room temperature. Sections were then incubated overnight at 4 °C with the indicated primary antibodies diluted in 3–5% BSA. The following day, sections were washed three times with PBS for 5 min each and incubated with appropriate secondary antibodies diluted in 3–5% BSA for 1–2 h at room temperature. Sections were washed again three times with PBS, mounted using Vectashield antifade mounting medium containing DAPI (Vector Laboratories), and imaged using a Nikon Ti Eclipse confocal microscope equipped with a ×40 oil-immersion objective. Image acquisition and analysis were performed in the Confocal and Specialized Microscopy Shared Resource of the Herbert Irving Comprehensive Cancer Center at Columbia University, supported by NIH/NCI Cancer Center Support Grant P30CA013696. For retinal morphological analysis, enucleated eyes fixed in formalin were embedded in paraffin. Sections (10 μm) were deparaffinized in xylene (3–5 min), rinsed in absolute ethanol (3–5 min), and rehydrated through graded ethanol solutions. Sections were stained with hematoxylin and eosin (H&E) and imaged using a Leica SCN400 whole-slide digital imaging system. Images were analyzed using Aperio ImageScope software (Leica). To detect lipid accumulation, paraformaldehyde-fixed sections were incubated with Nile Red (5 μg ml⁻¹ in 75% glycerol) for 30 min, rinsed with distilled water, and imaged by confocal microscopy. For RPE whole-mount staining, eye cups containing RPE and choroid were dissected and fixed in 4% paraformaldehyde for 1 h at room temperature, followed by washing with PBS. Eye cups were partially cut with four radial incisions and blocked with 5% BSA in PBST (0.3% Triton X-100). RPE/choroid flat mounts were incubated with phalloidin for 1 h and with anti-Iba1 antibody for 24 h, washed three times with PBS, and imaged using a Leica DM5000-B fluorescence microscope.

### Immunoblotting

RPE/choroid were isolated from WT, *Lamp2a^-/-^* untreated, and GLA1-1-treated *Lamp2a^-/-^* mice. Each sample was homogenized in 100 μl RIPA buffer, followed by sonication and centrifugation at 12,000 g for 20 min at 4°C. The resulting supernatant was collected for protein analysis. Cultured cells were lysed using CelLytic M cell lysis buffer (Sigma, C2978) on ice for 20 min and clarified by centrifugation. Protein concentrations were determined using the BCA protein assay (Thermo Fisher Scientific). Equal amounts of protein were separated by SDS–PAGE using NuPAGE Bis-Tris 4–12% gels (Novex, Life Technologies) and transferred to polyvinylidene difluoride (PVDF) membranes (Novex, Life Technologies). Membranes were blocked in StartingBlock blocking buffer (Thermo Fisher Scientific, 37543) for 1 h at room temperature and incubated overnight at 4°C with the indicated primary antibodies. The following day, membranes were washed three times for 5 min in TBST (0.2% Tween-20) and incubated with IR- or Alexa Fluor–conjugated secondary antibodies for 1 h at room temperature. Membranes were then washed three times with TBST and scanned using an Odyssey SA imaging system (LI-COR Biosciences, Lincoln, NE). Band intensities were quantified by densitometric analysis using ImageJ software.

### Transmission electron microscopy (TEM)

Enucleated eye cups were fixed overnight at 4°C in 0.1 M cacodylate buffer containing 2.5% glutaraldehyde and 2% paraformaldehyde. Fixed samples were dehydrated through a graded series of cold acetone and embedded in epoxy resin (Oken Epok 812; Oken Shoji, Tokyo, Japan). Ultrathin silver-gold sections were cut using a diamond knife, mounted on 50-mesh copper grids coated with a Formvar membrane, and stained with uranyl acetate and lead citrate. Sections were examined using a Hitachi HT7700 transmission electron microscope.

### Statistical analysis

Data are presented as mean ± SD. For animal experiments, n represents the number of mice analyzed in each group. For cell culture experiments, results are representative of at least three independent biological replicates. Statistical analyses were performed using GraphPad Prism 10. Differences among groups were assessed using one-way or two-way analysis of variance (ANOVA), as appropriate, followed by Tukey’s or Dunnett’s multiple-comparison tests. Exact statistical tests and sample sizes are indicated in the corresponding figure legends. Statistical significance was defined as *P < 0.05.

## Acknowledgments

This project was supported by the Edward N. and Della L. Thome Memorial Foundation Award in Age-Related Macular Degeneration Research (to KP and CLC), NIH R21EY035369 grant (to KP and CLC), P30 EY019007 (Core Support for Vision Research), and unrestricted funds from Research to Prevent Blindness (New York, NY) to the Department of Ophthalmology, Columbia University.

