## Supplementary Materials for "Loss of Lamp2a-dependent chaperone-mediated autophagy drives dry AMD-like retinal pathology in mice and is rescued by BK channel activation"

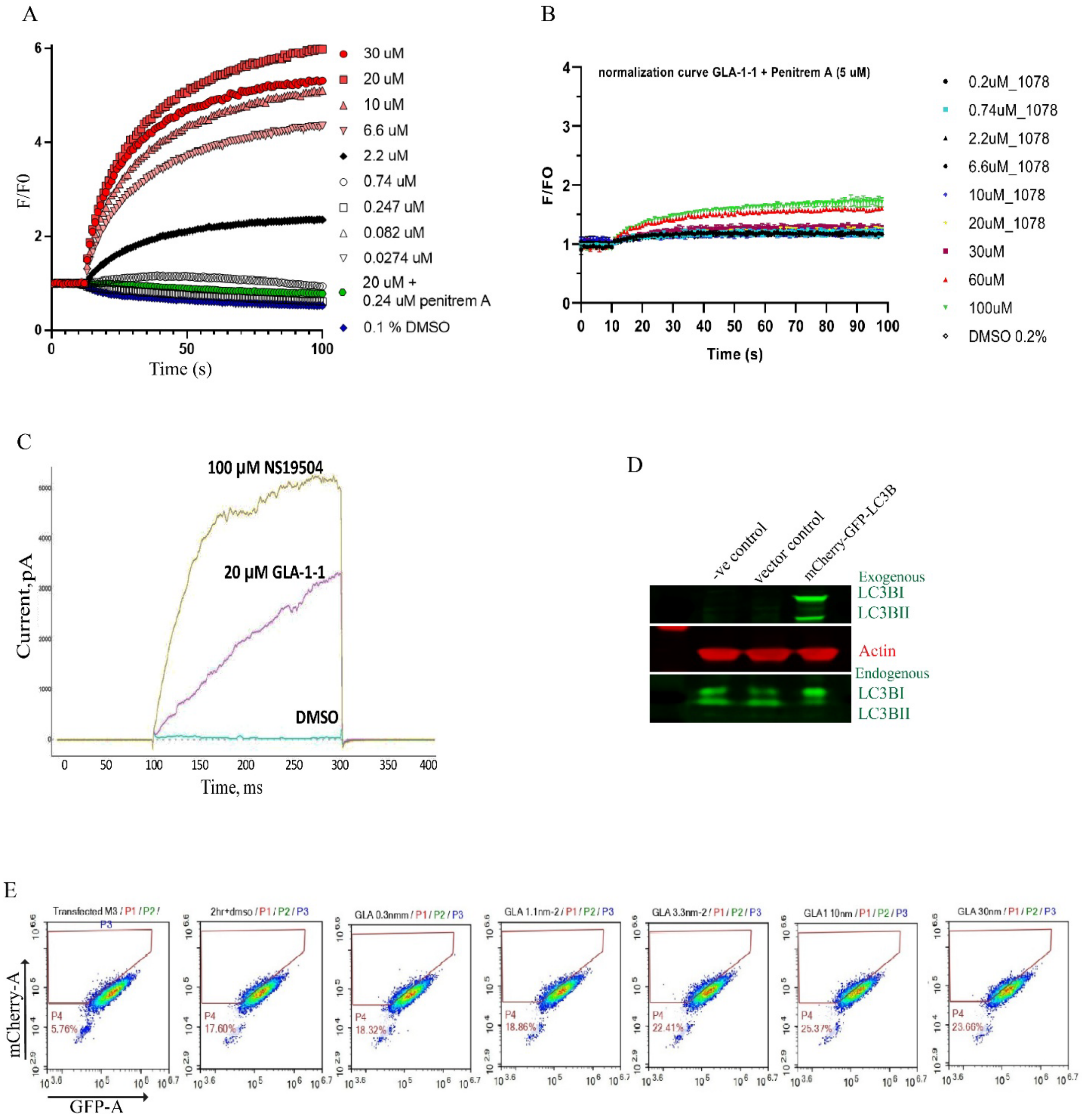


**Fig. S1 (supplementary): Representative figures of BK channel activation induced by BK agonists.** (A) Representative FLIPR traces showing BK channel activation by GLA-1-1 across a range of concentrations (0.0274–30 μM). The assay reports normalized fluorescence changes ($F/F_0$) reflecting membrane potential changes associated with BK channel activation. Fluorescence responses were recorded for 100 s following compound addition. (B) FLIPR traces showing the effect of the BK channel inhibitor penitrem A (5 μM) on GLA-1-1–induced responses. Co-treatment with penitrem A markedly suppresses the increase in normalized fluorescence ($F/F_0$), confirming that the signal observed in panel A is mediated by BK channel activation. Recording duration was 100 s. (C) Whole-cell patch-clamp recordings demonstrating BK channel currents induced by GLA-1-1 compared with the high concentration of the reference BK agonist NS19504. GLA-1-1 induces robust outward currents relative to DMSO control. (D) Immunoblot confirming expression of the tandem mCherry-EGFP-LC3B reporter following retroviral transduction. Exogenous LC3B expression is detected in transduced cells in addition to endogenous LC3B. Actin was used as a loading control. (E) Representative flow cytometry plots showing autophagic flux in HeLa cells expressing the mCherry-EGFP-LC3B reporter following treatment with increasing concentrations of GLA-1-1 (0.3–30 nM). The shift in mCherry/GFP signal indicates increased autophagic flux, with maximal activation observed at 10 nM (n = 2 technical replicates, n = 5 biological replicates).

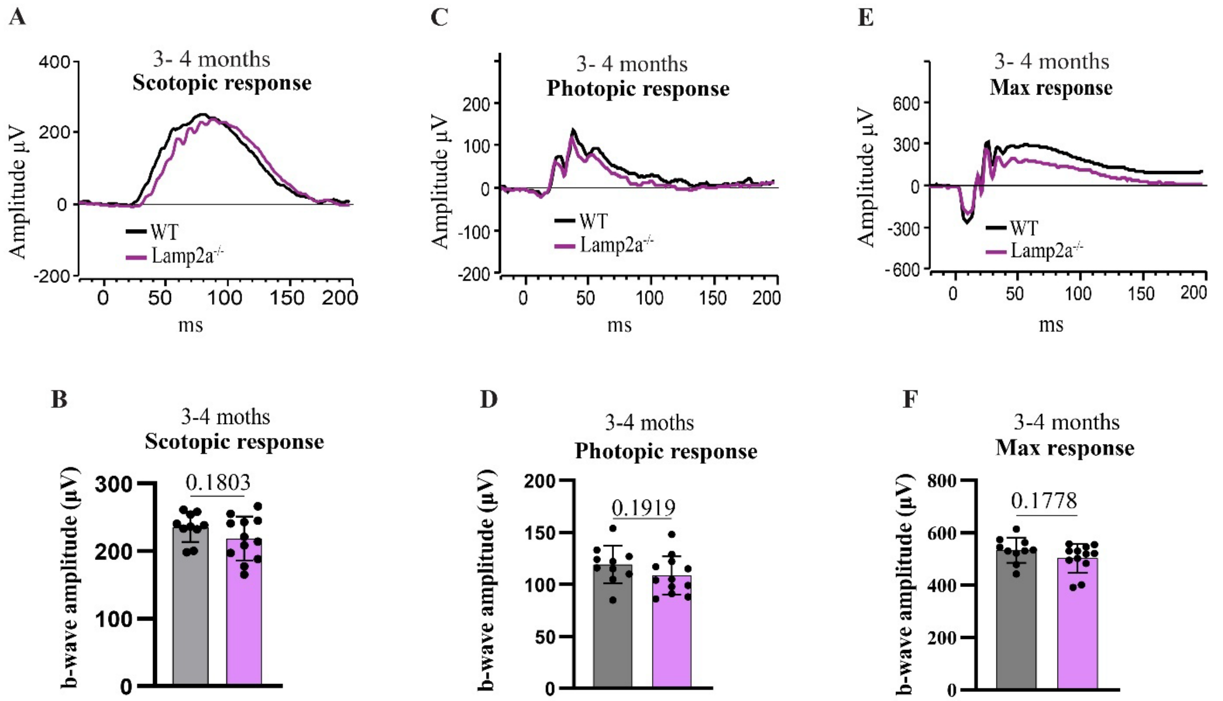


**Fig. S2 (supplementary): Electroretinography (ERG) analysis of retinal function in 3–4-month-old WT and *Lamp2a*$^{-/-}$ mice. (A)** Representative scotopic ERG traces recorded from dark-adapted WT and *Lamp2a*$^{-/-}$ mice in response to dim flash stimulation (0.001 cd·s·m$^{-2}$), reflecting rod-mediated responses. (**B**) Quantification of scotopic ERG b-wave amplitudes showing no significant difference between WT and *Lamp2a*$^{-/-}$ mice (WT, n = 10; *Lamp2a*$^{-/-}$ n = 12). **(C)** Representative photopic ERG traces recorded under light-adapted conditions (30 cd·s·m$^{-2}$ flash with rod-suppressing background illumination), reflecting cone-mediated responses. (**D**) Quantification of photopic ERG b-wave amplitudes showing no significant difference between WT and *Lamp2a*$^{-/-}$ mice (WT, n = 10; Lamp2a$^{-/-}$, n = 12). **(E)** Representative ERG traces showing the maximal mixed rod–cone response elicited by a bright flash stimulus (3 cd·s·m$^{-2}$) in dark-adapted mice. (**F**) Quantification of maximal mixed rod–cone b-wave amplitudes showing no significant difference between WT and *Lamp2a*$^{-/-}$ mice (WT, n = 10; Lamp2a$^{-/-}$, n = 12). Data are presented as mean ± SD. Statistical analysis was performed using an unpaired two-tailed Student's t-test.

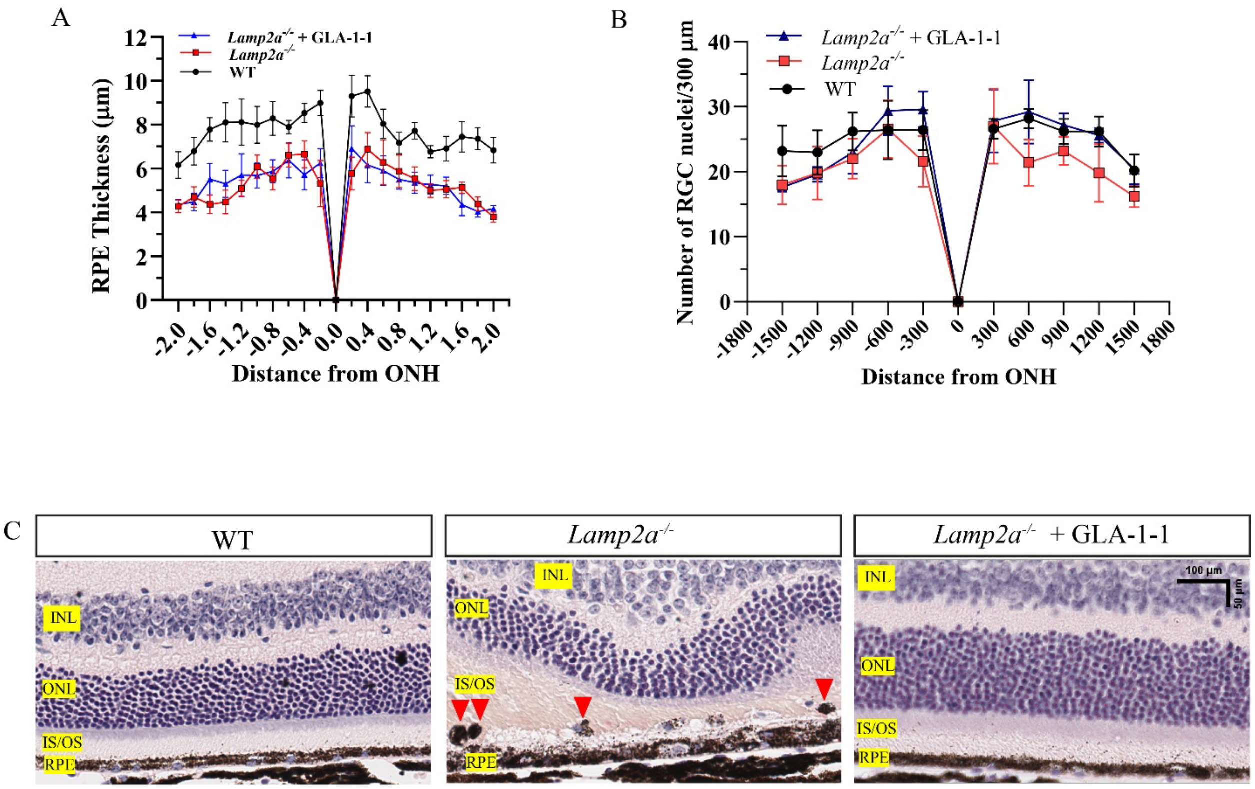


**Fig. S3 (supplementary): GLA-1-1 preserves RPE thickness, RGC counts, and RPE positioning in *Lamp2a$^{-/-}$* mice.** (**A**) Radial analysis of RPE thickness plotted as a function of distance from the optic nerve head (ONH) (n = 5 mice). *Lamp2a$^{-/-}$* mice showed reduced RPE thickness compared with WT mice, which was partially restored by GLA-1-1 treatment at multiple retinal locations. (**B**) Radial analysis of RGC nuclei counts plotted as a function of distance from the ONH (n = 5 mice). *Lamp2a$^{-/-}$* mice showed reduced RGC nuclei counts compared with WT mice; RGC nuclei count was restored by GLA-1-1 treatment. (**C**) Representative H&E-stained retinal sections showing anterior displacement of dysmorphic RPE cells toward the IS/OS region in *Lamp2a$^{-/-}$* mice. This abnormal RPE migration was not observed in GLA-1-1–treated *Lamp2a$^{-/-}$* mice.

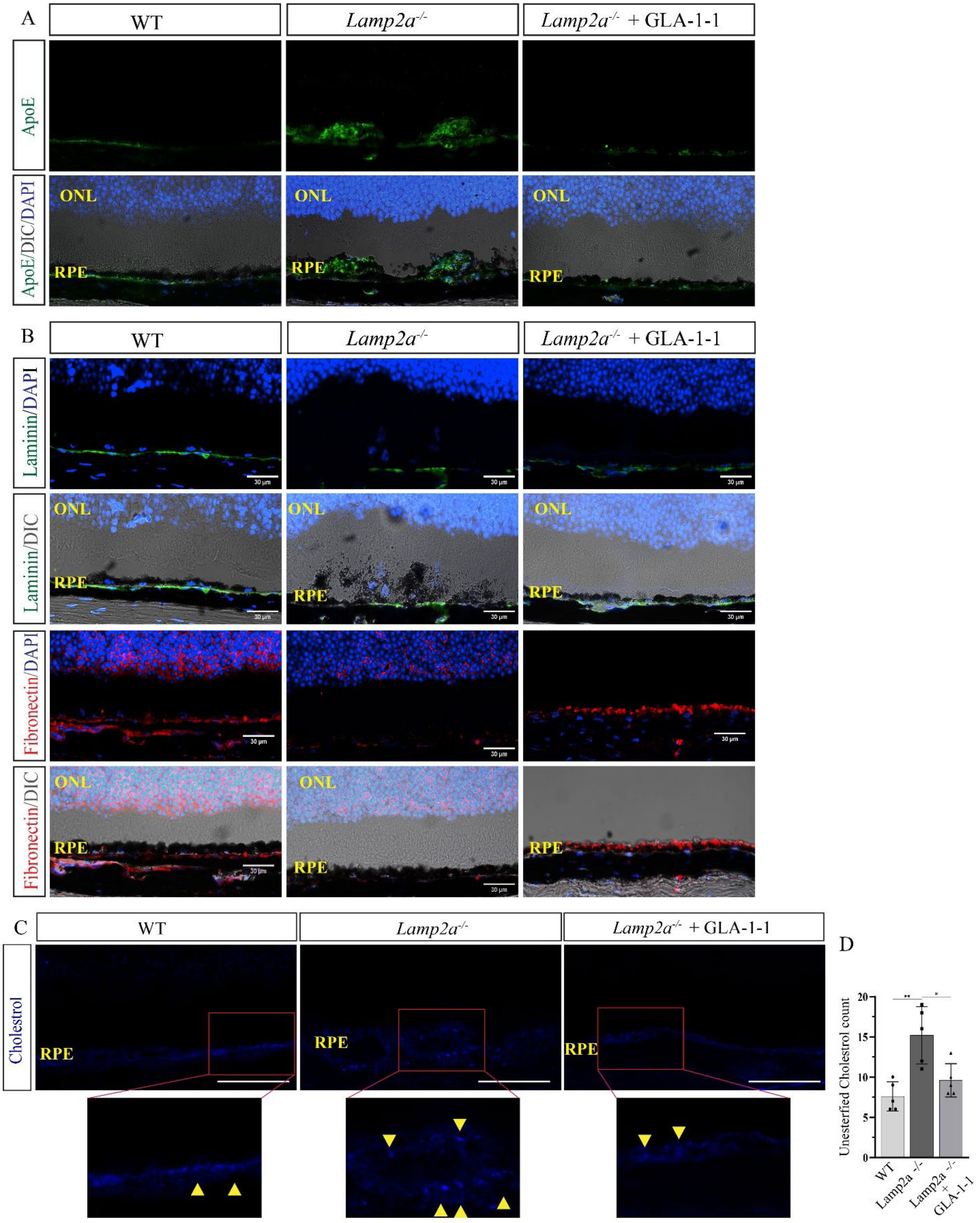


**Fig. S4 (supplementary): Lamp2a deficiency is associated with subretinal ApoE accumulation, extracellular matrix alterations, and increased unesterified cholesterol.** (**A**) Representative confocal immunofluorescence images showing ApoE staining (green) in retinal sections. *Lamp2a*$^{-/-}$ mice exhibited ApoE accumulation extending into the subretinal space above the RPE, which was reduced by GLA-1-1 treatment. DAPI, blue. Scale bar = 30 µm. (**B**) Representative confocal immunofluorescence images showing laminin (green) and fibronectin (red) staining in retinal sections. *Lamp2a*$^{-/-}$ mice showed reduced laminin and fibronectin immunoreactivity relative to WT mice, which was restored by GLA-1-1 treatment. DAPI, blue. Scale bar = 30 µm. (**C**) Representative filipin staining of retinal sections showing unesterified cholesterol in the RPE region. Insets show enlarged views of the boxed areas. Arrowheads indicate filipin-positive signal. Scale bar = 100 µm. (n=5 mice). (**D**) Quantification of filipin-positive unesterified cholesterol signal showing increased levels in *Lamp2a*$^{-/-}$ mice compared with WT mice, which were reduced by GLA-1-1 treatment (n = 5 mice). Statistical analysis was performed using one-way ANOVA with Tukey's post hoc test. $^{*}P < 0.05$, $^{**}P < 0.01$. Values are expressed as mean ± SD.

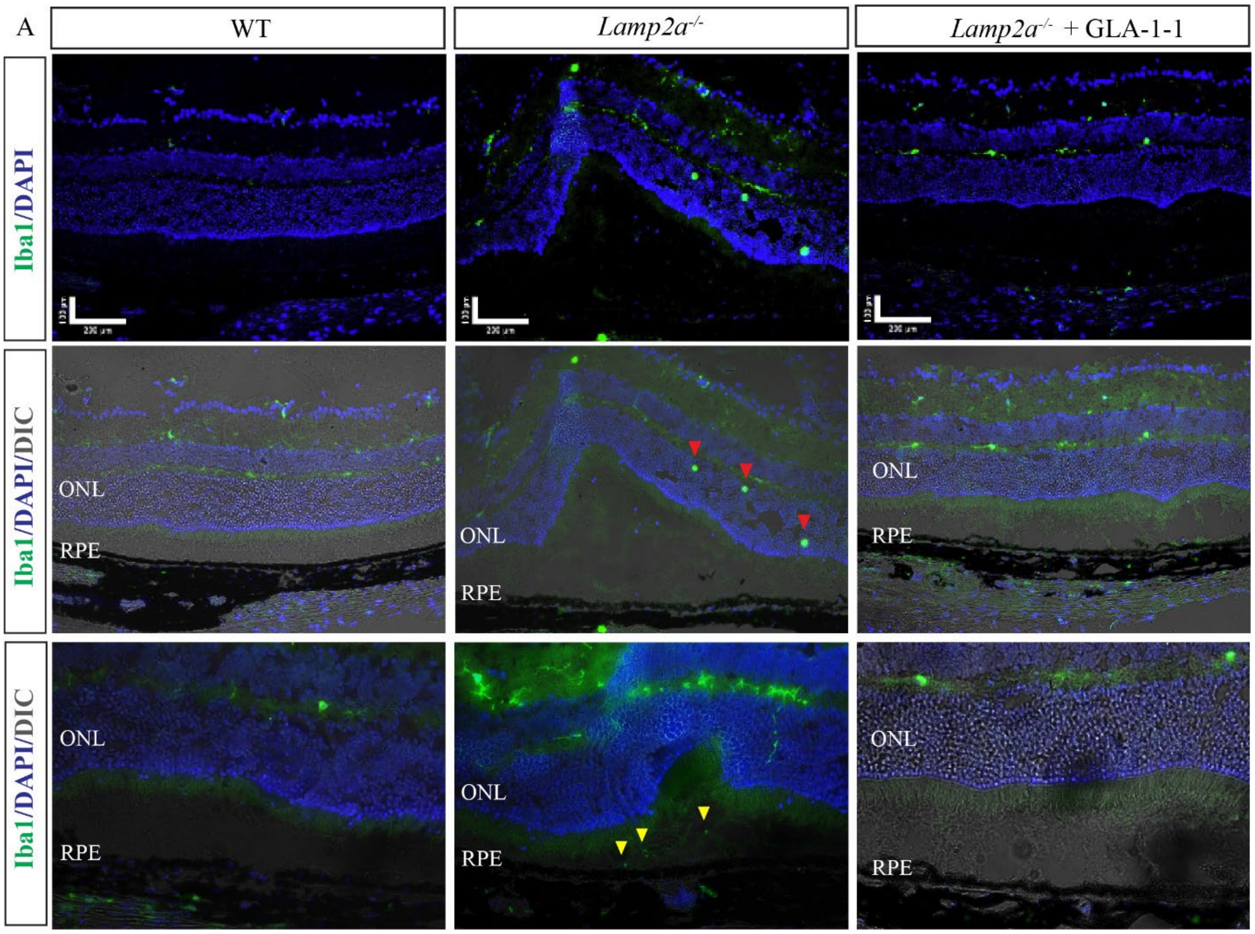


**Fig. S5 (supplementary): Lamp2a deficiency increases Iba1⁺ cell accumulation and subretinal migration.** Representative Iba1 immunostaining of retinal cross sections showing increased numbers of Iba1⁺ microglia/macrophages in *Lamp2a*$^{-/-}$ mice relative to WT mice. Activated Iba1⁺ cells with amoeboid morphology (red arrowheads) were observed migrating toward the photoreceptor IS/OS region, and additional Iba1⁺ cells were present in the subretinal space (yellow arrowheads). These abnormalities were reduced by treatment with the BK channel agonist GLA-1-1.

Table S1

Intracellular GLA-1-1 concentrations in mCherry-EGFP-LC3B reporter HeLa cells following incubation with GLA-1-1 at 10 nM and 30 nM.

| Analyte | Sample ID | GLA-1-1 concentration in culture media | Intracellular concentration, μM |
|---|---|---|---|
| GLA-1-1 | 001-GLA-1-1-10nM | 10 nM | 0.95 |
| | 002-GLA-1-1-10nM | | 1.15 |
| | 003-GLA-1-1-10nM | | 0.97 |
| | 004-GLA-1-1-30nM | 30 nM | 3.01 |
| | 005-GLA-1-1-30nM | | 5.06 |
| | 006-GLA-1-1-30nM | | 2.82 |

Table S2

PCR primers used for genotyping Lamp2a knockout mice

| Primer sequence |
|---|
| F1: 5'-CAACTACATAAACAGCATGAAGCCA-3'<br>R1: 5'-TCTTGGAGAAGCAAGAGCATAACT-3'<br>Mutant allele: 392 bp; Wildtype allele: 1947 bp |
| F2: 5'-CAGGCTCAGAAAGTTATAACAGAGC-3'<br>R1: 5'-TCTTGGAGAAGCAAGAGCATAACT-3'<br>Mutant allele: no amplification; Wildtype allele: 500 bp |

Table S3

Serum concentrations of GLA-1-1 measured in compound-treated mice.

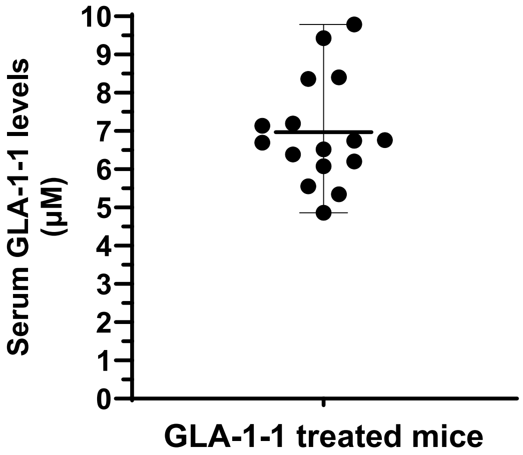
